# An aging clock using metabolomic CSF

**DOI:** 10.1101/2021.04.04.438397

**Authors:** Nathan Hwangbo, Xinyu Zhang, Daniel Raftery, Haiwei Gu, Shu-Ching Hu, Thomas J. Montine, Joseph F. Quinn, Kathryn A. Chung, Amie L. Hiller, Dongfang Wang, Qiang Fei, Lisa Bettcher, Cyrus P. Zabetian, Elaine Peskind, Gail Li, Daniel E.L. Promislow, Alexander Franks

**Affiliations:** Department of Statistics and Applied Probability, University of California, Santa Barbara, Santa Barbara, CA; Northwest Metabolomics Research Center, Department of Anesthesiology and Pain Medicine, University of Washington School of Medicine, Seattle WA; Veterans Affairs Puget Sound Health Care System, Seattle, WA; Department of Neurology, University of Washington School of Medicine, Seattle, WA; Department of Pathology, Stanford University School of Medicine, Palo Alto, CA; Portland Veterans Affairs Medical Center, Portland, OR; Department of Neurology, Oregon Health and Science University, Portland, OR; Department of Psychiatry and Behavioral Sciences, University of Washington School of Medicine, Seattle, WA; Department of Biology, University of Washington, Seattle WA; Department of Lab Medicine & Pathology, University of Washington School of Medicine, Seattle WA

**Author notes:** Corresponding author: Alexander Franks.

**Keywords:** metabolomics, cerebrospinal fluid, aging clock, biomarker

## Abstract

Quantifying the physiology of aging is essential for improving our understanding of age-related disease and the heterogeneity of healthy aging. Recent studies have shown that in regression models using “-omic” platforms to predict chronological age, residual variation in predicted age is correlated with health outcomes, and suggest that these “omic clocks” provide measures of biological age. This paper presents predictive models for age using metabolomic profiles of cerebrospinal fluid from healthy human subjects, and finds that metabolite and lipid data are generally able to predict chronological age within 10 years. We use these models to predict the age of a cohort of subjects with Alzheimer’s and Parkinson’s disease and find an increase in prediction error, potentially indicating that the relationship between the metabolome and chronological age differs with these diseases. In our analysis of control subjects, we find the carnitine shuttle, sucrose, biopterin, vitamin E metabolism, tryptophan, and tyrosine to be the most associated with age. We showcase the potential usefulness of age prediction models in a small dataset (n = 85), and discuss techniques for drift correction, missing data imputation, and regularized regression which can be used to help mitigate the statistical challenges that commonly arise in this setting. To our knowledge, this work presents the first multivariate predictive metabolomic and lipidomic models for age using mass spectrometry analysis of cerebrospinal fluid.

## 1 Introduction

Risk of age-related disease varies among individuals, and is shaped by genetic factors, environmental factors, and the interaction between the two(1). As with most complex traits, in attempts to map specific genetic variants that are associated with phenotypic variation, researchers have identified genes that are significantly correlated with age-related disease, but which explain only a small fraction of the overall variation(2),(3), contributing to the so-called “missing heritability” problem(4). In order to address the large functional distance between genotype and phenotype, researchers have turned to ‘endophenotypes’–transcriptome, epigenome, metabolome, lipidome, and microbiome–as a way to bridge the gap between genotype and phenotype, and characterize the physiological processes of aging(5).

In our own recent work, we have focused on the metabolome, which measures the structural and functional building blocks of an organism, as a powerful link between genotype and phenotype in studies of aging and age-related traits(6),(7). During this same period, there has been considerable interest in the power of the epigenome to explain variation in patterns of aging within human populations. These prior studies have established predictive models for age using epigenetic data(8),(9), and these “epigenetic clocks” have been shown to be useful biomarkers for risk factors of mortality(10). While chronological age of a cohort of people all born, say, in 1961, is the same (60 yrs in 2021), the underlying biological age varies. Aging research has focused on markers of underlying physiological age, that is, as a “biological clock”.

Here we focus on the degree to which variation in the metabolome, specifically as measured in cerebrospinal fluid (CSF), might function as a “metabolomic clock”. Researchers have found that variation in the metabolome, which measures small molecules (< 2000 Da) circulating in an organism, can account for variation in diverse traits, including all-cause mortality(11),(12). Furthermore, individual and small sets of metabolite concentrations have been found to be associated with age(13),(14). Rather than focus on the association between a single metabolite and age, we use complete metabolomic profiles to create predictive models for age. Since the metabolome is the endophenotype furthest downstream in the genotype-phenotype path, a predictive model for age using the metabolome might provide unique insight into the physiological mechanisms of aging.

Metabolomic clocks using urine, serum, and plasma samples have been found to be predictive of chronological age(15)–(18). In the present study, we construct predictive models for age analyzing both targeted and untargeted metabolomic and lipidomic profiles of CSF. CSF serves to cushion the brain as well as transport biological substances, and is the fluid in closest proximity to the central nervous system, making it valuable for analysis, particularly with respect to age-related neurological disease(19). There is some evidence that the relationship between metabolites in the brain and age is distinct from other organs, suggesting that a predictive model using CSF might provide new insight into aging in the brain(20),(21).

The invasive nature of collecting CSF typically means that large sample sizes are difficult to obtain. To our knowledge, this is the first attempt at creating a predictive model for age using CSF data. This study is also notable for the comprehensive set of metabolomic profiles analyzed here, including targeted metabolomics, global metabolomics, and lipidomic profiles. Previous efforts to create biological clocks from high-dimensional data(8),(15) have relied on a class of statistical models known as *Elastic Net* regularized linear regression. Such models are commonly used to form these age-prediction models because of their ability to handle the case when the number of features (e.g. methylation sites, metabolites) greatly exceeds the number of samples, and to perform feature selection in the process(16),(22),(23). We discuss techniques and challenges involved in fitting these models in small sample, high-dimensional profiles, including methods for missing data imputation and cross-validation, as well as methods to mitigate bias that can occur when fitting these high dimensional regression models. We also address broader statistical challenges in these -omic based “biological clock” analyses, such as accounting for batch effects and signal intensity drift over time. We carry out pathway analysis and metabolite set enrichment analysis using univariate relationships between metabolites and age, and find that the carnitine shuttle, sucrose, biopterin, vitamin E metabolism, tryptophan, and tyrosine are associated with age. Finally, in addition to analyzing healthy individuals, we also use our models to assess any evidence of accelerated aging in the metabolome profiles of patients with Alzheimer’s disease (AD) and Parkinson’s disease (PD).

## 2 Materials and Methods

### 2.1 Biological samples

One hundred ninety-eight CSF samples, collected from 85 controls, 57 AD patients, and 56 PD patients were available for analysis. The 85 healthy subjects range from 20 to 86 years of age with a median of 56, while the AD and PD subjects’ range from 35 to 88 years of age with a median of 67. We also record each subject’s sex at birth. Data on *APOE* ε4 genotype (all participants) and *GBA* carrier status (pathogenic mutations and the E326K polymorphism for the PD group only) were generated in previous work by our group and made available for this study(24),(25).

For the control and AD subjects, CSF samples were provided from the VA Northwest Mental Illness Research, Education, and Clinical Center (MIRECC) Sample and Data Repository. Participants underwent neurological examination and a detailed neuropsychological assessment. Participants were determined to be cognitively normal controls or AD patients by expert clinical diagnosis confirmed by neuropsychological testing and Clinical Dementia Rating scale. All controls had a Clinical Dementia Rating scale score of 0, MMSE scores between 26-30, and paragraph recall scores no lower than < 1 SD below age- and education-matched norms. AD participants met NINDS-ADRDA criteria for probable AD.

PD CSF was obtained from participants enrolled in the Pacific Udall Center at the Veterans Affairs (VA) Puget Sound Health Care System and the VA Portland Health Care System. All PD subjects underwent a neurological examination and a detailed neuropsychological assessment. The resulting data were reviewed at a joint consensus conference to ensure diagnostic consistency between the two sites as previously described(26),(27). Only participants who met U.K. Parkinson’s Disease Society Brain Bank clinical diagnostic criteria for PD were included in this study(28).

At all sites CSF was collected in the fasting state in the morning. CSF was collected in the lateral decubitus position using the Sprotte 24g atraumatic spinal needle. CSF was collected by negative pressure into sterile polypropylene syringes and aliquoted into 0.5 ml aliquots in polypropylene cryotubes and frozen immediately on dry ice at the bedside. All samples were stored at −80° C prior to assay.

The study procedures were approved by the institutional review boards of the University of Washington, VA Puget Sound Health Care System, and VA Portland Health Care System. All participants, or their legally authorized representative for those with impaired decisional capacity, provided written informed consent.

### 2.2 Metabolomics

Samples were aliquoted into four different sub-samples, and subsequently prepared for four distinct metabolomic profiles, with three aqueous metabolite profiles, including i) targeted metabolomics; ii) global metabolomics; iii) Globally Optimal Targeted Mass Spectrometry (GOT-MS); and iv) lipidomics.

#### 2.2.1 Targeted metabolomics

Targeted metabolomics analysis was carried out using an LC-MS/MS platform targeting 203 standard metabolites from more than 25 metabolic pathways (e.g., glycolysis, tricyclic acid cycle, amino acid metabolism, glutathione, etc.). LC-MS/MS experiments were performed on a Waters Acquity I-Class UPLC TQS-micro MS system. Each sample was injected twice, 2 μL and 10 μL, for analysis using positive and negative ionization modes, respectively. Both chromatographic separations were performed in hydrophilic interaction chromatography mode. The flow rate was 0.3 mL min^−1^, autosampler temperature was kept at 4 °C, and the column compartment was set at 40 °C. The mobile phase was composed of solvents A (5 mM ammonium acetate in H2O + 0.5% acetic acid + 0.5% acetonitrile) and B (acetonitrile + 0.5% acetic acid + 0.5% water). The LC gradient conditions were the same for both positive and negative ionization modes. After an initial 1.5-min isocratic elution of 10% A, the percentage of solvent A was increased linearly to 65% at time (t) = 9 min, then remained the same for 5 min (t = 14 min), and then reduced to 10% at t = 15 min to prepare for the next injection. After chromatographic separation, MS ionization and data acquisition was performed using an electrospray ionization source. A pooled study sample was used as the quality control (QC) and run once for every 10 study samples.

#### 2.2.2 Global metabolomics

Global MS-based lipidomics was performed using an Agilent 1200 LC system coupled to an Agilent 6520 Q-TOF mass spectrometer. CSF samples (200 μL) were prepared by dissolving in 1000 μL methanol, which was vortexed, incubated at – 20 °C and centrifuged at 18000 x g for 20 min. Then. 750 μL were collected, dried and then reconstituted in a 100 μL solution of 40% H_2_O 60% acetonitrile. Five μL of each prepared sample were analyzed by positive ESI ionization and 10 μL analyzed by negative ESI ionization. Samples were separated using a Waters XBridge BEH Amide column (15 cm x 2.1mm, 2.5 μm), which was heated to 40 °C. The mobile phase was 10 mM NH_4_HCO_3_ in 100% H_2_O (Solvent A) and 100 % acetonitrile (Solvent B) and its gradient was 95-10% B from 0 to 5 min, 10% B from 5-40 min., 10-100% B from 40-45 min., and 100% from 40-70 min. MS parameters were performed according to previously described methods(29). The mass accuracy of our LC−MS system is generally better than 5 ppm, the Q-TOF/MS spectrometer was calibrated prior to each batch run, and a mass accuracy of <1 ppm was often achieved using the standard tuning mixture (G1969-85000, Agilent Technologies). The m/z scan range was 100−2000, and the acquisition rate was 1.0 spectra/s. MS data were processed using Agilent Mass Hunter and Mass Profiler Professional.

#### 2.2.3 GOT-MS

Globally Optimized Targeted MS (GOT-MS) is a technique which sits between targeted and untargeted metabolomic profiling(30),(31). The GOT-MS method used here was modeled after Zhong, Xu, and Xhu, 2019(32), and are detailed in the Supplemental eMethods.

#### 2.2.4 Lipidomics

Lipids were extracted from the samples (200 uL) using dichloromethane/methanol after the addition of 54 isotope labeled internal standards across 13 lipid classes. The extracts were concentrated under nitrogen and reconstituted in 100 uL solution consisting of 10 mM ammonium acetate in dichloromethane:methanol (50:50). Lipids were analyzed using the Sciex Lipidyzer platform consisting of a Shimadzu Nexera X2 LC-30AD pumps, a Shimadzu Nexera X2 SIL-30AC autosampler and an AB Sciex QTRAP 5500 MS/MS system equipped with SelexION for differential mobility spectrometry (DMS) according to the methods we developed previously(33). Multiple reaction monitoring (MRM) was used to target and quantify over 1000 lipids in positive and negative ionization modes with and without DMS. Data were acquired and processed using Sciex Analyst 1.6.3 and Lipidomics Workflow Manager 1.0.5.0. A key for lipid abbreviations is available in Supplemental eTable 1.

### 2.3 Data Analysis

#### 2.3.1 Data Pre-Processing

Samples were prepared for assay in seven batches of 28 or 29 samples each. To mitigate batch effects, we explicitly balanced age, sex, phenotype (controls, Alzhiemer’s, Parkinson’s), and APOE status across batches. To this end, we used a variant of the “finite selection model” proposed in Morris et al.(34). For each of the seven batches, we iteratively selected the subject that was the most different from the subjects already assigned to that batch, with respect to the aforementioned covariates. This process balances the samples across the batches, which is important for correcting drift and maximizing power to detect differences. Covariate balance across batch is shown in eFigure 1 in the Supplement.

Prior to inference, we corrected for systematic drift in the mass spectrometer inferred intensities over time. The observed drifts were metabolite-specific and thus separate corrections were done for each metabolite. In Supplemental eFigure 2,we show examples of metabolites for which the raw intensities showed significant changes in mean intensity over time, along with the data after correction. To account for the very general functional form of the intensity drift over time, we used extreme gradient boosting, a tree-based ensemble method that can recover non-continuous functions. We use the xgboost package in R and cross-validation to select tuning parameters and control over-fitting(35).

After drift correction, we centered and scaled each feature, and treated observations further than 3 Median Absolute Deviations (MAD) within each feature as missing values to be imputed. Features with more than 50% missingness were removed.

#### 2.3.2 Missing Data Imputation

We use the R package Amelia to estimate the missing values in our dataset(36). This method creates multiple completed versions of the dataset under the assumption that the data are well approximated by a multivariate normal distribution, and that the missing values are Missing at Random (MAR), which is to say, the differences between the missing and observed values can be fully explained using the observed variables in the dataset. We created five imputed datasets of each profile and ran our analysis on each of these completed datasets. The variation across imputed datasets provides a measure of robustness to the imputation procedure. In Figure 3,the error bars indicate the minimum, maximum, and average result among the five imputations.

Because this method requires that the number of subjects exceeds the number of metabolites, we assume that each metabolite follows a normal distribution, so that the underlying distribution of the transposed data is multivariate normal. In cases of large amounts of missingness, we add a ridge prior, shrinking the assumed covariances between metabolites. These modifications make it possible to complete the imputation at cheaper computational cost, but add additional bias to the imputed values.

#### 2.3.3 Principal Component Analysis (PCA) and Partial Least Squares-Discriminant Analysis (PLS-DA)

Principal Component Analysis (PCA) is a technique commonly used for visualizing high dimensional data by projecting it onto a lower dimensional space. This unsupervised procedure identifies dimensions that maximize the amount of preserved variability in the data. In addition to PCA, we can also make use of a supervised method known as Partial Least Squares. Partial Least Squares maximizes the covariance between the data and a given variable. Partial Least Squares - Discriminant Analysis (PLS-DA) is the special case where the provided variable is categorical. Both PCA and PLS-DA were implemented using the NIPALS algorithm in the R package ropls, which allows for missing values(37). We combine these two techniques in the following way:

- Perform PCA on the combined profiles dataset *X* to obtain a loading matrix *W* and score matrix *T*, which satisfies *X* = *TW*^*T*^.
- Extract the first column of *T* (*t*_1_) and the first row of 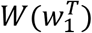 and compute 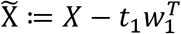.
- Run PLS-DA on 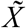 with respect to sex and plot the first PLS component against the first principal component from PCA (*t*_1_).

#### 2.3.4 Regression Modeling

Gaussian elastic net regression models extend the linear regression model to allow for the case where the number of features exceeds the number of observations, and can be implemented using the glmnet package in R(22). Two parameters need to be specified by the user to implement this model: *α* and *λ*. Following Horvath, 2013(8), we use *α* = 0.5, which performs variable selection but avoids randomly choosing one out of a set of correlated predictors (a problem with models fit with *α* = 1). *λ*, the overall penalty parameter, is chosen to minimize squared error in leave one out cross-validation (LOOCV).

The only metadata included in the model is the sex of each subject. Because of the putative sex-related differences in the metabolome(16),(38), we assume sex at birth is *a priori* an important predictor and do not perform shrinkage for this variable.

To evaluate the performance of these models, we proceed in a leave-one-out fashion by iteratively omitting a single observation and training a model on the remaining data. For each observation, we proceed as follows:

1. Remove features missing from the left-out observation from the training set, following the reduced modeling methodology validated in Saar-Tsechansky and Provost, 2007(39).
2. Impute the remaining missing values from the training set to create five imputed versions of the dataset.
3. Fit models on the five imputed training sets, using LOOCV to tune *λ*
4. Predict the age of the left-out observation in each imputation.

Applying this process to every observation and comparing the predictions to chronological age yields the plots seen in Figure 3. To obtain a snapshot of the features driving these predictions, a model is fit on all the control subjects. The coefficients for this full model in the targeted and lipidomic profiles are reported in Table 1. The full model on the untargeted profile is then used to predict the age of an out-of-sample cohort containing 57 AD and 56 PD subjects. These patients tended to be older and in a smaller age range than our healthy cohort. Thus, rather than comparing the predictions for the AD/PD group using error metrics for the full set of controls, we opt for the following procedure:

1. For each subject in the new cohort, find the control with the same sex and closest age. If there are ties, then pick the subject with the closest processing batch number.
2. Using the untargeted model fit on the full set of controls, compute a predicted age for each of the AD/PD subjects.
3. Compare results between the two sets.

**Table 1:**
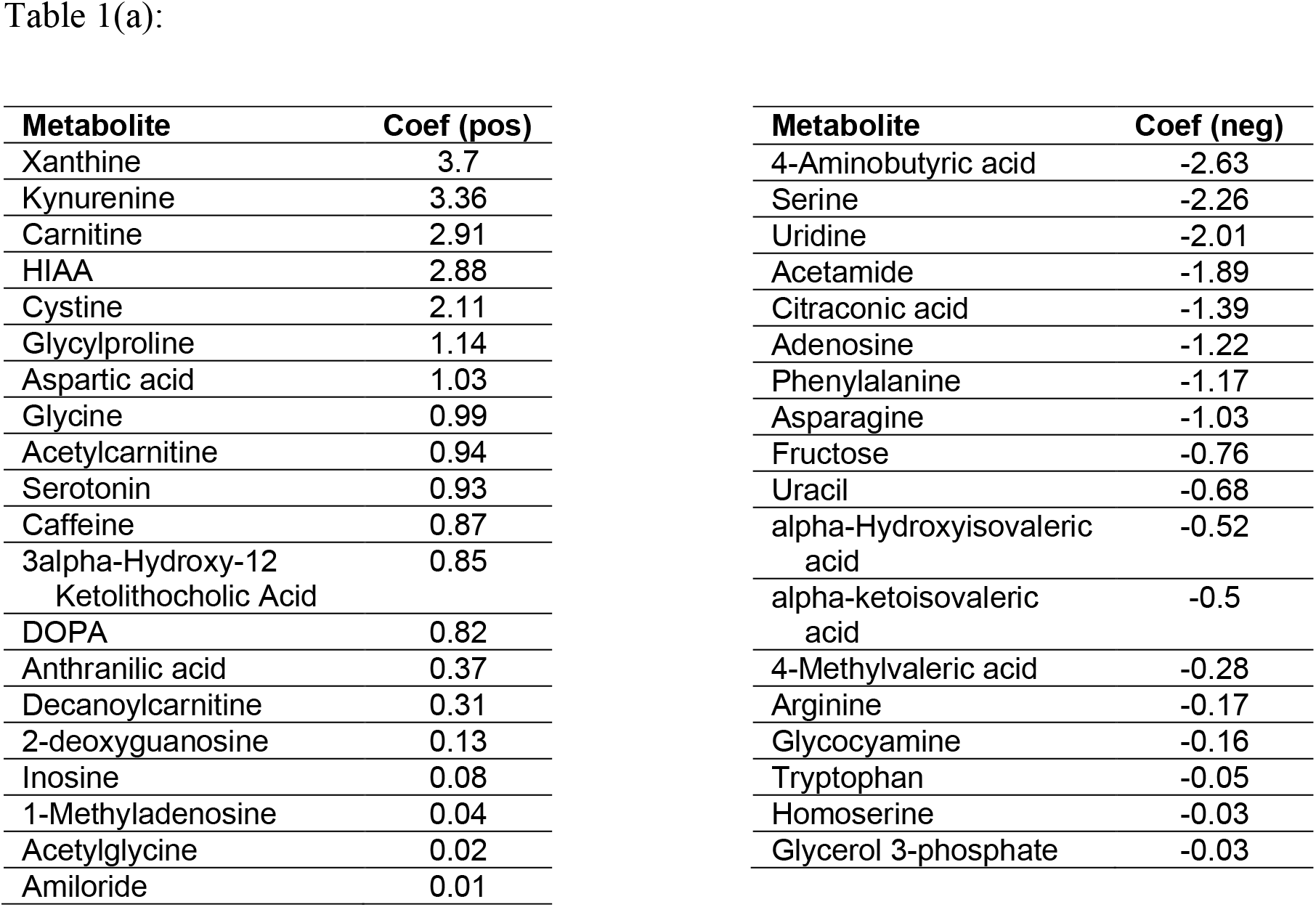
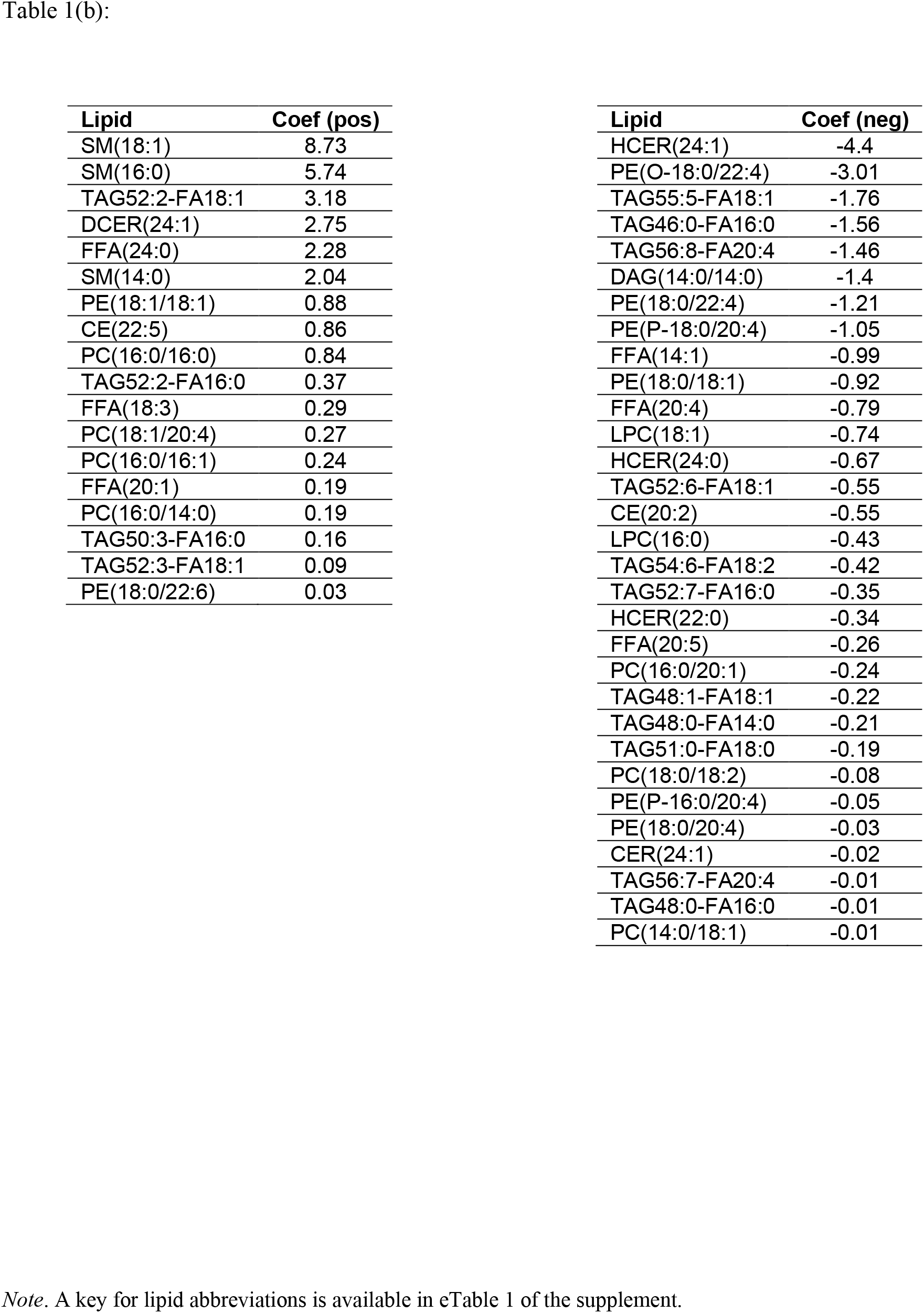
Names and average coefficient for targeted metabolites (a) and lipids (b) retained in elastic net models fit on the full data. The tables are sorted by magnitude and split into positive and negative coefficient tables. Sex at birth was not penalized, and is therefore on a different scale from the coefficients in the tables. With this in mind, being male led to a decreased age prediction of 0.90 years in the targeted profile, and a decreased age prediction of 2.87 years in the lipids profile.

| Metabolite | Coef (pos) | Metabolite | Coef (neg) |
| --- | --- | --- | --- |
| Xanthine | 3.7 | 4-Aminobutyric acid | -2.63 |
| Kynurenine | 3.36 | Serine | -2.26 |
| Carnitine | 2.91 | Uridine | -2.01 |
| HIAA | 2.88 | Acetamide | -1.89 |
| Cystine | 2.11 | Citraconic acid | -1.39 |
| Glycylproline | 1.14 | Adenosine | -1.22 |
| Aspartic acid | 1.03 | Phenylalanine | -1.17 |
| Glycine | 0.99 | Asparagine | -1.03 |
| Acetylcarnitine | 0.94 | Fructose | -0.76 |
| Serotonin | 0.93 | Uracil | -0.68 |
| Caffeine | 0.87 | alpha-Hydroxyisovaleric acid | -0.52 |
| 3alpha-Hydroxy-12 Ketolithocholic Acid | 0.85 | alpha-ketoisovaleric acid | -0.5 |
| DOPA | 0.82 | 4-Methylvaleric acid | -0.28 |
| Anthranilic acid | 0.37 | Arginine | -0.17 |
| Decanoylcarnitine | 0.31 | Glycocyamine | -0.16 |
| 2-deoxyguanosine | 0.13 | Tryptophan | -0.05 |
| Inosine | 0.08 | Homoserine | -0.03 |
| 1-Methyladenosine | 0.04 | Glycerol 3-phosphate | -0.03 |
| Acetylglycine | 0.02 |  |  |
| Amiloride | 0.01 |  |  |

Table 1(b):
| **Lipid** | **Coef (pos)** |
| --- | --- |
| SM(18:1) | 8.73 |
| SM(16:0) | 5.74 |
| TAG52:2-FA18:1 | 3.18 |
| DCER(24:1) | 2.75 |
| FFA(24:0) | 2.28 |
| SM(14:0) | 2.04 |
| PE(18:1/18:1) | 0.88 |
| CE(22:5) | 0.86 |
| PC(16:0/16:0) | 0.84 |
| TAG52:2-FA16:0 | 0.37 |
| FFA(18:3) | 0.29 |
| PC(18:1/20:4) | 0.27 |
| PC(16:0/16:1) | 0.24 |
| FFA(20:1) | 0.19 |
| PC(16:0/14:0) | 0.19 |
| TAG50:3-FA16:0 | 0.16 |
| TAG52:3-FA18:1 | 0.09 |
| PE(18:0/22:6) | 0.03 |

| **Lipid** | **Coef (neg)** |
| --- | --- |
| HCER(24:1) | -4.4 |
| PE(O-18:0/22:4) | -3.01 |
| TAG55:5-FA18:1 | -1.76 |
| TAG46:0-FA16:0 | -1.56 |
| TAG56:8-FA20:4 | -1.46 |
| DAG(14:0/14:0) | -1.4 |
| PE(18:0/22:4) | -1.21 |
| PE(P-18:0/20:4) | -1.05 |
| FFA(14:1) | -0.99 |
| PE(18:0/18:1) | -0.92 |
| FFA(20:4) | -0.79 |
| LPC(18:1) | -0.74 |
| HCER(24:0) | -0.67 |
| TAG52:6-FA18:1 | -0.55 |
| CE(20:2) | -0.55 |
| LPC(16:0) | -0.43 |
| TAG54:6-FA18:2 | -0.42 |
| TAG52:7-FA16:0 | -0.35 |
| HCER(22:0) | -0.34 |
| FFA(20:5) | -0.26 |
| PC(16:0/20:1) | -0.24 |
| TAG48:1-FA18:1 | -0.22 |
| TAG48:0-FA14:0 | -0.21 |
| TAG51:0-FA18:0 | -0.19 |
| PC(18:0/18:2) | -0.08 |
| PE(P-16:0/20:4) | -0.05 |
| PE(18:0/20:4) | -0.03 |
| CER(24:1) | -0.02 |
| TAG56:7-FA20:4 | -0.01 |
| TAG48:0-FA16:0 | -0.01 |
| PC(14:0/18:1) | -0.01 |

*Note.* A key for lipid abbreviations is available in eTable 1 of the supplement.

### 2.4 Pathway Analysis

Mummichog is a tool used for pathway analysis of untargeted metabolomic data, accessible via a script using the Python programming language(40). To use this program, we input mass:charge ratio and retention time for each feature, as well as *t*-values and (Benjamini-Hochberg corrected) *p*-values taken from a univariate linear model regressing age on each feature. The output is a comparison of the input with known metabolomic pathways, split by the features’ mode.

For pathway analysis on the targeted profile, Metabolite Set Enrichment Analysis (MSEA) is a tool which finds associations between sets of metabolites by performing a hypergeometric test(41). MSEA is implemented using the MetaboAnalystR package, which takes as input the names of significant metabolites (using a false discovery rate (FDR) < 0.05 from a univariate linear regression), as well as the names of all the features in our dataset to be used as a reference set(42).

## 3 Results

The samples were processed using untargeted and targeted metabolite profiling, as well as Globally Optimized Targeted Mass Spectrometry (GOT-MS) and lipid profiling. The untargeted, targeted, GOT-MS, and Lipid profiling yielded 6735, 108, 854, and 1070 features, respectively. These samples came with widely varying degrees of missingness, with 16%, 3%, 4%, and 81% of the data missing from each of the profiles, respectively. Supplemental eFigure 3 displays a flowchart outlining the analysis performed on each of the four profiles.

### 3.1 PCA/PLS-DA

PCA on the dataset formed by combining all four profiles showed that the first two principal components explain 20% of the variance in our dataset. Subjects with negative scores for the first principal component have a median age of 70, while subjects with a positive score have a median age of 28, indicating that the largest source of variation in the profiles is driven by subject age. We also apply PLS-DA (a supervised method) on the combined dataset in order to simultaneously separate the profiles by sex. We find that these first two PLS components account for only 11% of the variation in our data. Following the procedure described in Section 2.3.3, Figure 2 shows the combined data projected onto the first principal component from PCA (PC1), along with the first PLS component discriminating sex on the orthogonal complement of PC1.

**Figure 1:**
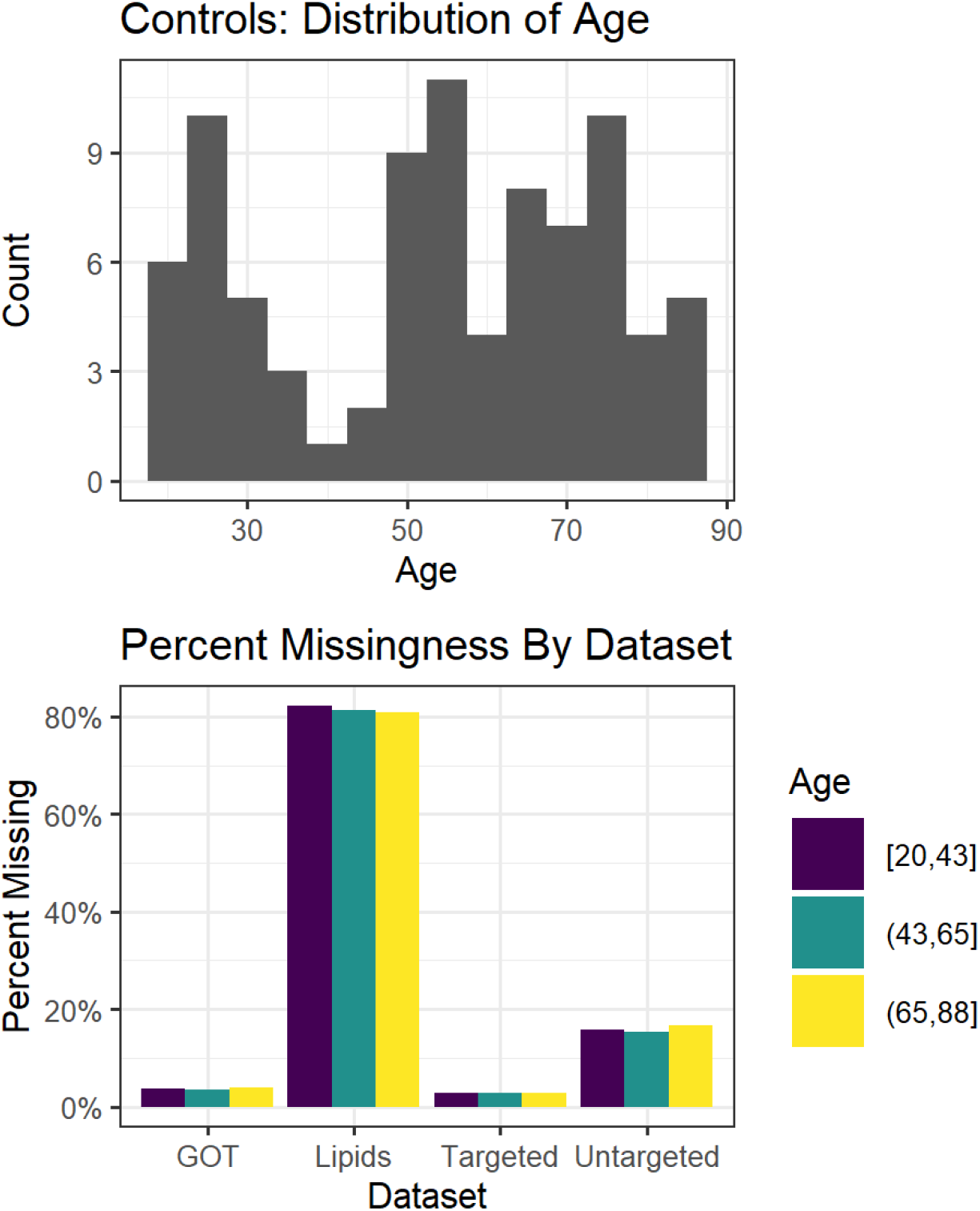
Distribution of age (top) as well as a comparison of missingness between profiles, split in three approximately equal sized age groupings (bottom)

**Figure 2:**
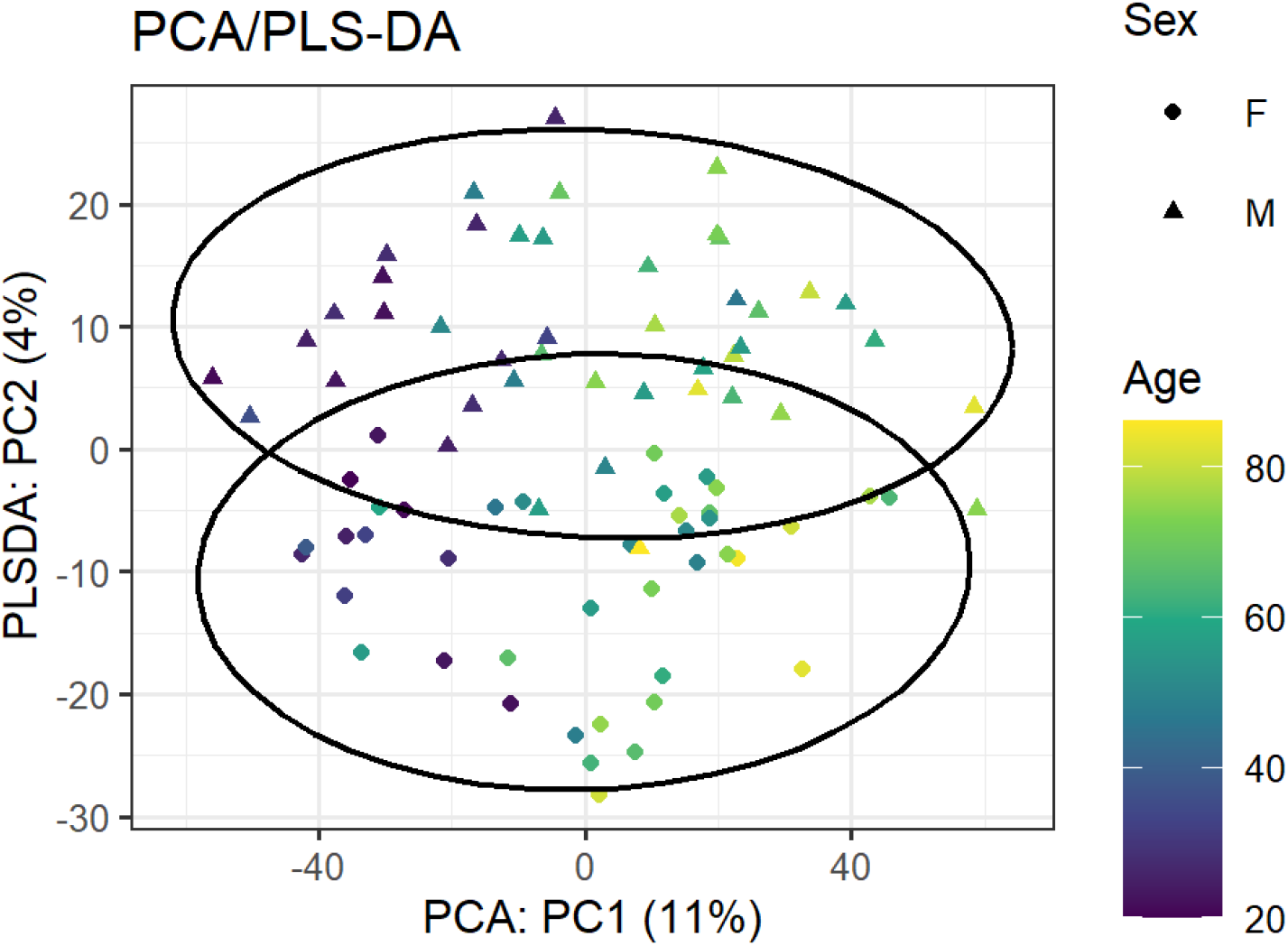
Separation of sex and age using the combined result of PCA and PLS-DA on the dataset of combined profiles. The x-axis displays the first principal component from PCA, while the y-axis displays the first PLS component discriminating sex at birth on the space orthogonal to x-axis. For PLS-DA, the first two principal components are plotted, with confidence ellipses (assuming *t* distribution). In parentheses, the axis names contain the percent variation (of the dataset) explained by that component.

### 3.2 Missing Data

While investigating the varying degrees of missingness in our data, we found that the youngest patients tended to have the largest amount of missing data across all the datasets, with the most missingness in the lipidomic data. To quantify the relationship between age and missingness on aggregate, we fit an elastic net regression model where we replace metabolite concentrations with 0 if the value is missing, and 1 otherwise. For this model, we report a Root Mean Squared Error (RMSE) of 14.8, Mean Absolute Error (MAE) of 12.4 and an R^2^ of 0.47 in the untargeted dataset (Supplemental eFigure 4).

To characterize the relationship between age and missingness for individual lipids, we separate subjects into three equally spaced age groups, and perform Pearson Chi-Squared tests on each lipid, testing the null hypothesis that the proportion of missingness is the same within each cohort. These tests found six lipids with FDR < 0.05, all of which are triglycerides (Supplemental eFigure 4), which represents a 2.2-fold enrichment over what would be expected if six lipids were selected by chance. More broadly, our results demonstrate the importance of examining differential missingness across groups in the ‘omics of aging.

### 3.3 Predictive Models for Age

We train separate age-prediction models on each of our four datasets, and found a MAE ranging from ∼ 7 − 13 years, with the lowest error from the untargeted dataset. Figure 3 depicts chronological age against the predicted age for each of these profiles, while Supplemental eFigure 5 displays the same figure for the model combining all four profiles. The predictions using each of the four profiles (after detrending for chronological age) report Spearman rank-order correlations between 0.1 and 0.54, with the predictions from the lipids profile reporting the lowest correlations (Supplemental eTable 1). As a reference point, the intercept-only model, which predicts the mean age for every subject, yields an RMSE of 20.18 and MAE of 17.18.

**Figure 3:**
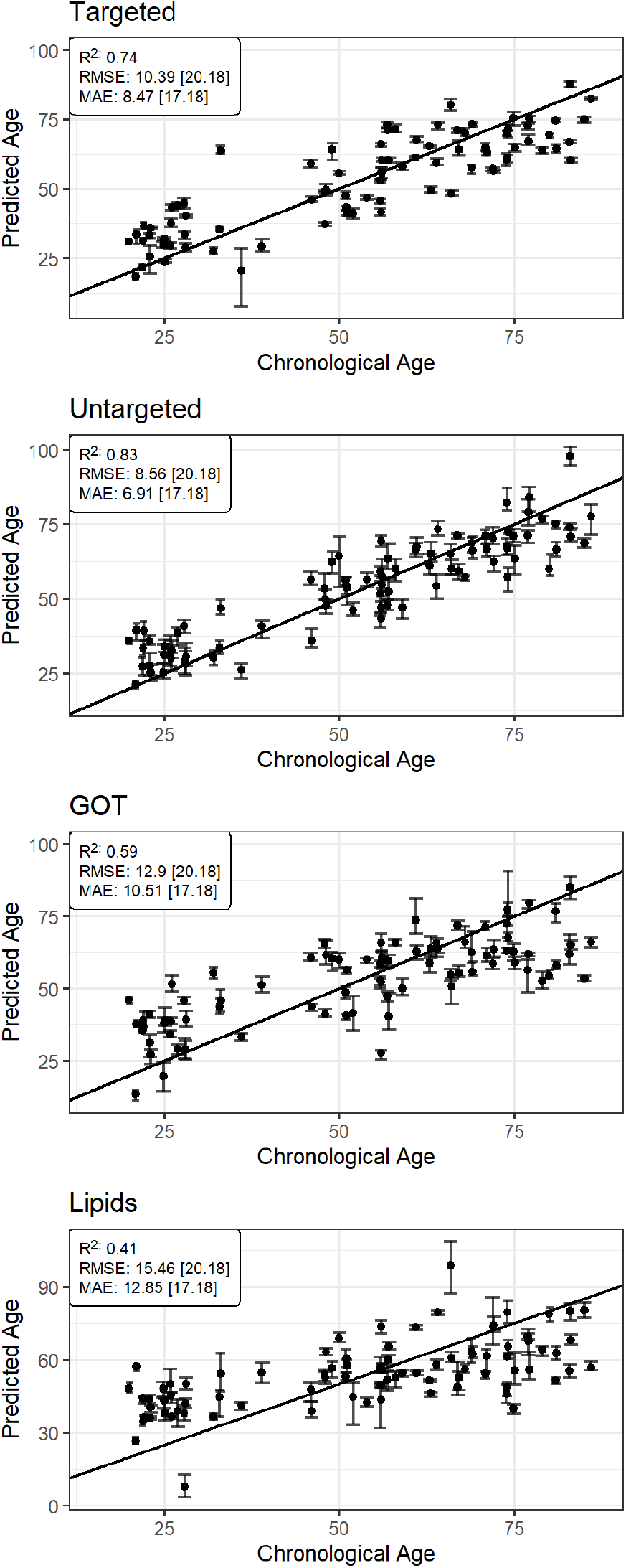
Predicted vs. chronological Age for controls in each profile, with R^2^, Root Mean Squared Error (RMSE), and Mean Absolute Error (MAE) reported. Numbers in parentheses represent the performance of the mean model for comparison. Points are the average of the predictions for the five imputations to estimate missing values, with the error bars representing the most extreme predicted values from the imputations. We also include the *y* = *x* line. Points above the line correspond to overestimates of a subject’s chronological age, while points below the line correspond to underestimates.

Additionally, models are fit on three variations of the untargeted profile to test the effects of drift correction in data preprocessing, to verify the modeling decision to keep sex exempt from penalization, and to test a biological age hypothesis. First, we verify that the drift correction procedure described in Section 2.3.1 improves out-of-sample predictions for age compared to using the uncorrected metabolite intensities in the untargeted profile, reporting an RMSE of 10.3, MAE of 8.4, and R^2^ of 0.75. Compared to the performance of the models fit after drift correction (Figure 3), this represents an 18% increase in RMSE, 20% increase in MAE, and 10% reduction in R^2^.

To verify the decision to exclude sex from regularization, models for males and females were fit separately on the untargeted profile. We obtain RMSEs of 11.1 and 12.8, MAEs of 9.5 and 9.8, and R^2^ of 0.7 and 0.6, for males and females respectively. There are no metabolites consistently shared between the separate sex models among the five missing data imputed datasets, possibly indicating that the separate sex models are picking up different signal in the metabolome. Performing the same procedure on random samples of controls of the same sizes as the separate sex models (44 for males and 41 for females), we obtain similar predictive accuracy: RMSEs of 10.7 and 12.64, MAEs of 8.7 and 10.3, and R^2^ of 0.68 and 0.54, for males and females respectively.

A previous epigenetic clock study based on 353 CpG sites tends to over-predict the age in a cohort of subjects with Parkinson’s disease(43). To test whether our metabolomic age prediction model exhibits similar behavior, we obtained age predictions for a cohort of 57 AD subjects and 56 PD subjects. Figure 4 displays the predictions of an age-matched control cohort alongside the AD/PD cohort for comparison. We find that the predictions for the AD/PD cohort are less accurate than those for the age-matched controls, with a 50% decrease in R^2^, 30% increase in RMSE, and 18% increase in MAE. We also observe that within the Parkinson’s cohort, the subject with the largest model error was one of the six pathogenic GBA carriers in our dataset.

**Figure 4:**
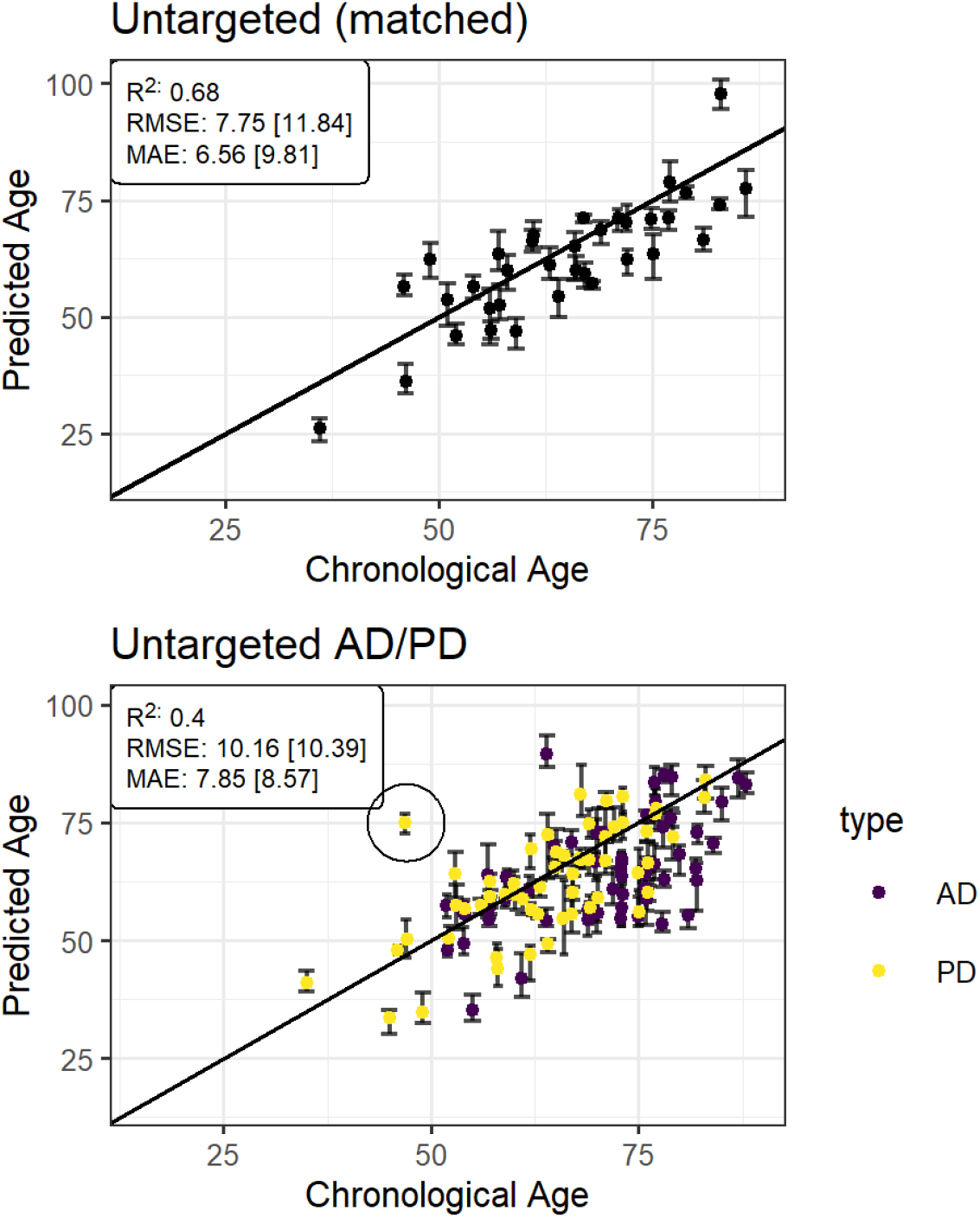
The leave one out performance of the untargeted model only looking at matched controls (left) and the performance of the model on the AD/PD cohort (right). The circled point is the greatest outlier (in terms of RMSE) for PD. Numbers in parentheses refer to the performance of the model which predicts the mean age for each subject.

### 3.4 Age-associated Metabolites

Elastic net automatically performs variable shrinkage by setting coefficients for non-predictive metabolites to zero. Our models for the untargeted, targeted, and GOT-MS datasets typically included non-zero coefficients for between thirty and forty metabolites. Supplemental eFigure 6 shows summary statistics for the behavior of these models across the imputations and leave one out modeling procedure.

The models applied to the lipidomic data tended to include fewer features than the metabolite datasets, with an average of 15 retained across the five imputations. Table 1 lists the targeted metabolites and lipids that appeared in models fit on all 85 control subjects. As a summary, we find that in the model formed using the targeted profile, increased concentrations of xanthine, kynurenine, carnitine, HIAA, and cystine were the largest contributors to higher age predictions, while increased concentrations of 4-aminobutyric acid, serine, and uridine were the largest contributors to lower age predictions. In the lipidome model, increased concentrations of SM(18:1), SM(16:0), TAG52:2-FA 18:1, DCER(24:1), FFA(24:0), and SM(14:0) were the largest contributors to higher age predictions, while increased concentrations of HCER(24:1) and PE(O18:0/22:4) were the largest contributors to lower age predictions. Because the features were standardized prior to fitting the models, the listed coefficients represent the expected difference in age prediction resulting from a one standard deviation increase in concentration. Note, however, that because our models are multivariate, these coefficients represent change in expected age given all of the other covariates, and should therefore not be interpreted unconditionally.

### 3.5 Pathway Analysis and Set Enrichment Analysis

While the untargeted dataset yielded the smallest predictive error among all sets, the metabolite identities are unknown, making biological interpretation difficult. At FDR < 0.05, Mummichog identified the carnitine shuttle, starch and sucrose metabolism, putative anti-inflammatory metabolites formation from EPA, and biopterin metabolism pathways for the positive mode, and the glyoxylate and dicarboxylate metabolism pathway for the negative mode as associated with age (Supplemental eFigure 7).

We also use the targeted data to identify metabolomic pathways with an overrepresentation of metabolites that are significantly associated with age by MSEA(41). Using the Small Molecule Pathways Database (SMPDB)(44) or MetaboAnalyst’s CSF disease-associated library, MSEA finds no associations below an FDR threshold of 0.05. However, our list of significant metabolites has large overlap with the tryptophan and tyrosine metabolism set in the SMPDB (Supplemental eTable 3).

## 4 Discussion

Two seminal papers authored by Horvath(8) and Hannum(9) on epigenetic clocks inspired investigation into an assortment of biological clocks(5), including the metabolomic clock proposed by Rist et al., 2017(16) and Robinson et al., 2020(15). In this paper, we present predictive models for age using CSF metabolomic data. We find that metabolite and lipid data are generally able to predict chronological age within approximately 10 years. For the profiles where feature identity is known (targeted metabolomics and lipidomics), the model coefficients are reported to indicate which predictors are driving the model. To our knowledge, this work represents the first metabolomic and lipidomic clock using mass spectrometry analysis of CSF.

### 4.1 Biological Interpretation of Targeted Multivariate Analysis

In the age prediction model built on the targeted metabolomic profile, we find that several of the metabolites driving the predictions (listed in Table 1) have been found to be associated with age. Consistent with our results, Blennow et al., 1993(45) finds a positive correlation between 5-hydroxy-indoleacetic acid (5-HIAA) and age in human CSF, and Johnson et al., 2019(14) reports negative correlation between serine and age in human plasma.

In the age prediction model fit on the lipidomic profile, sphingomyelins (SM) are the largest drivers of age predictions, as three appear with positive associations with age. Collino et al., 2013(46) finds that SM(16:0) concentration increased with age (although other SM species exhibited the opposite behavior).

### 4.2 Biological Interpretation of Pathways Analysis

The pathways identified by Mummichog from the untargeted profile are consistent with the existing literature on age-related changes in the metabolome. The carnitine shuttle has previously been identified to be associated with age in humans(13). Komori et al., 1999(47) finds a relationship between the biopterin metabolism and age using CSF samples. The vitamin E metabolism has been reported to be associated with age by Robinson et al., 2020(15) in a metabolomic aging analysis. In a study of mice, Ivanisevic et al., 2016(21) identify purine to be related with aging in the brain. Several of these pathways, the carnitine shuttle, Vitamin E metabolism and tryptophan metabolism observed here have also been associated with frailty (48).

MSEA does not find any significant associations for targeted univariate analysis at FDR < 0.05. However, the *p*-values for this procedure are computed using Over-Representation Analysis based on the cumulative hypergeometric distribution to compare the input list of significant metabolites to the SMPDB. As such, our input list can contain all the metabolites in small SMPDB pathways (in our case, tyrosine and glutamate metabolism) and the FDR would still be above 0.05. For our exploratory purposes, these associations can still be valuable. In particular, tryptophan has been featured prominently in the literature on the aging metabolome(7),(46), and association between tyrosine and age has been found in females in a longitudinal metabolomic study of humans(49).

### 4.3 Interpreting Aging Models Across Profiles

We observed considerable differences in results between profiles. These are likely caused by a combination of true differences between the datasets, regularization bias, and varying amounts of preprocessing needed to run the models. For instance, the lipids model was most sensitive to the effects of missing data, as 80% of the dataset was missing. Many features were excluded (those with *>* 50% missingness), and the rest were affected by missing data imputation. This lends itself to regularization bias, as the amount of signal in a feature is dampened due to the imputed values. Supplemental eDiscussion 1 showcases two attempts at reducing regularization bias in the modeling procedure: pre-selecting features using a univariate correlation threshold, and post-selection by elastic net. Neither method significantly improved results, but we find that post-selection inference can be useful when the researcher expects a small number of features to explain a large percentage of the variation in age (although it is likely invalid in the small data setting), while feature reduction by univariate correlation can be useful when the researcher expects the predictive power of the metabolites to be largely independent of one another. The comparable errors between these models suggests the presence of many equally performant age prediction models of the metabolome and can possibly explain the similar performance of the separate male and female models despite having no consistently shared metabolites.

Consideration should also be made for computational efficiency. An empirical prior is used to speed up the missing data imputation process but comes at the cost of shrinking covariances between predictors. This tradeoff is necessary because of the complexity of the leave-one-out prediction procedure, which is used to approximate out-of-sample performance in a small data setting. A separate validation set would have eliminated the need for these simplifications altogether.

We also note that our data represent a cross-sectional sample, which comes with inherent limitations when studying a temporal variable. As such, the pathways identified by the univariate analysis of the untargeted and targeted profiles are not attempts to claim understanding of the biological mechanisms behind aging. Rather, they are most helpful when considered in the context of existing work, and as a starting point for more focused future work.

### 4.4 Alzheimer’s, Parkinson’s and the biological clock

In the application of our age-prediction models to the AD/PD subjects, the performance is not much better than the intercept-only model, and there is no evidence of the consistent over-prediction present in Horvath and Rist, 2015(43). On the contrary, our models exhibit a pattern of under-prediction on these subjects. This behavior could be due to differences in the metabolome of the AD/PD cohort which cannot be attributed to age. However, this behavior could also be unrelated to disease status, as a pattern of underestimating the age of older subjects appears in the age-matched control cohort in Figure 4,and is also a pattern observed in epigenetic clocks(50). This under-prediction could be due to regularization bias induced by our modeling procedure, but it is also possible that metabolite concentrations become less informative of age as time progresses. Supplemental eFigure 11 flips our regression around, regressing individual metabolite concentrations by age, and demonstrating a possible leveling off of metabolite concentration at older ages.

A limitation to this analysis is that the models used to generate predictions for the AD/PD cohort could not be properly validated on control subjects. This is because we opted to form new elastic net models using all 85 of the control subjects, rather than applying each of the leave one out prediction models. However, the model coefficients are similar enough to the leave-one-out models to expect the predictive performance to be on par with the predictions shown in Figure 3.

### 4.5 Conclusion

A metabolomic analysis of the CSF has the potential to capture physiological variation that changes slowly over time, and that reflects variation in the central nervous system. However, it is important to keep in mind that the methods and results are limited due to the small size and cross-sectional nature of our data, which makes the “metabolomic age” established in this paper less reliable for use as an aging biomarker, as metrics akin to epigenetic “age acceleration” are impacted by these constraints(43). However, these limitations can be common when working with data from such invasive procedures as the extraction of CSF samples. We hope that this paper serves to illustrate methods that can be used to extract maximal information in these small sample size, large feature datasets, as well as showcase the predictive power and usefulness of such studies. We also hope that this work will motivate larger studies and analysis of longitudinal cohorts, with the goal of developing a more robust aging model using the metabolome.

## Funding

This work was supported by the National Institutes of Health (grant numbers P50 NS062684, P50 AG05136, S10 OD021562); the Department of Veterans Affairs (grant number 101 CX001702); the Veterans Affairs Northwest Mental Illness Research, Education, and Clinical Center, and an anonymous foundation. DELP was supported by the National Institutes of Health (grant numbers R01 AG049494, R01 AG057330). DELP and AF were supported by the National Institutes of Health (grant number R03 CA211160). Metabolomic assays were supported in part by the UW Nathan Shock Center of Excellent for the Biology of Aging grant P30 AG013280.

## Supplemental Material

### eMethods: Globally optimized Targeted Metabolomics

All LC-MS/MS experiments were performed on a Waters Acquity I-Class UPLC TQS-micro MS system (Milford, MA). Each 100 *μ*L CSF sample was prepared by extraction with methanol, then dried and reconstituted using 90:10 water/acetomitrile. Each sample was injected twice; 10 *μ*L for analysis using negative ionization mode and 4 *μ*L for analysis using positive ionization mode. Both chromatographic separations were performed in hydrophilic interaction chromatography (HILIC) mode on a Waters XBridge BEH Amide column (150 × 2.1 mm, 2.5 μm particle size, Waters Corporation, Milford, MA). The flow rate was 0.3 mL/min, auto-sampler temperature was kept at 4° C, and the column compartment was set at 40°C. The mobile phase was composed of Solvents A (5 mM ammonium acetate in 90% H2O/10% acetonitrile/0.2% acetic acid) and B (5 mM ammonium acetate in 90% acetonitrile/10% H2O/0.2% acetic acid). After the initial 1 min isocratic elution of 90% B, the percentage of Solvent B decreased to 40% at *t* = 11 min. The composition of Solvent B maintained at 40% for 4 min (*t* = 15 min), and then the percentage of B was gradually increased back to 90%, to prepare for the next injection. The mass spectrometer is equipped with an electrospray ionization (ESI) source. Targeted data acquisition was performed in multiple-reaction-monitoring (MRM) mode. We monitored 595 GOT-MS precursor ions and 1890 MRM transitions, under positive and negative ionization modes in the mass range of 60-600 Dal. The whole LC-MS system was controlled by Waters MassLynx software (Waters, Milford, MA). The extracted MRM peaks were integrated using the Waters TargetLynx software (Waters, Milford, MA).

**eTable 1:**
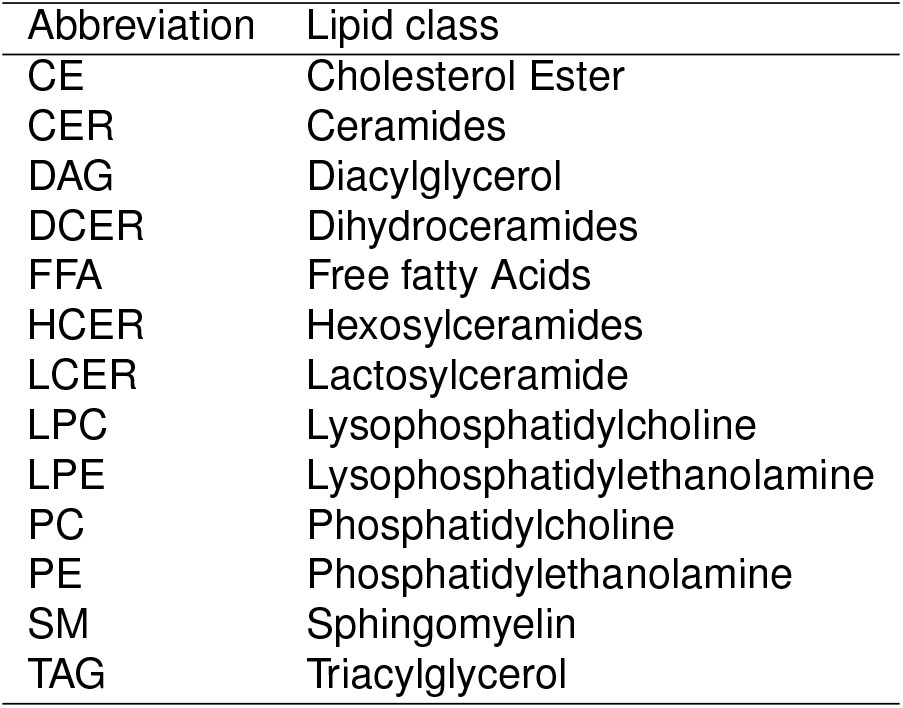
Key for Lipids Abbreviations.

| Abbreviation | Lipid class |
| --- | --- |
| CE | Cholesterol Ester |
| CER | Ceramides |
| DAG | Diacylglycerol |
| DCER | Dihydroceramides |
| FFA | Free fatty Acids |
| HCER | Hexosylceramides |
| LCER | Lactosylceramide |
| LPC | Lysophosphatidylcholine |
| LPE | Lysophosphatidylethanolamine |
| PC | Phosphatidylcholine |
| PE | Phosphatidylethanolamine |
| SM | Sphingomyelin |
| TAG | Triacylglycerol |

**eFigure 1:**
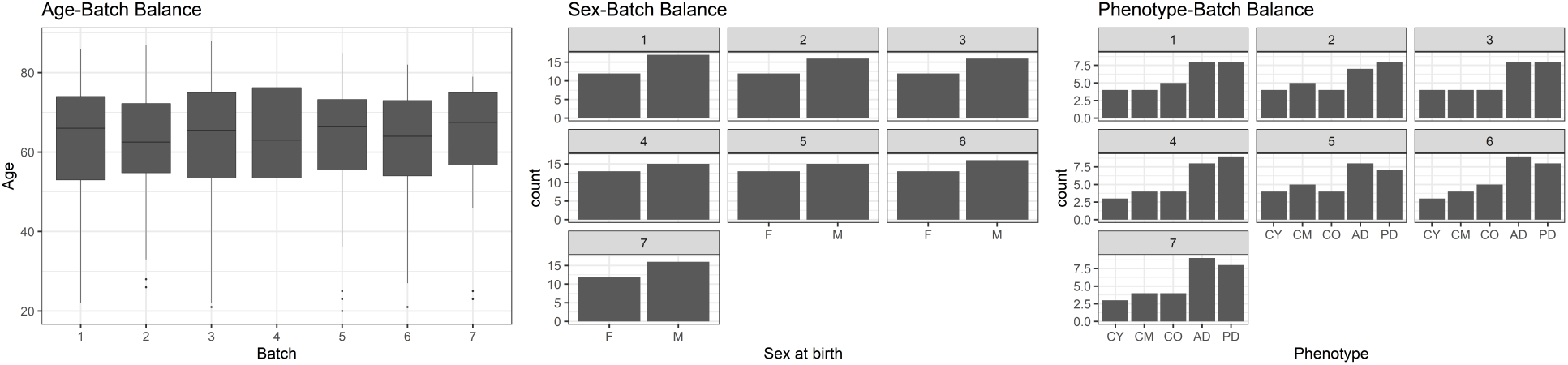
Balance across batches. Balance of age, sex, and phenotype across batches. This lessens the effect batch number could have in our predictive models, making the models more likely to pick up true biological differences.

**eFigure 2:**
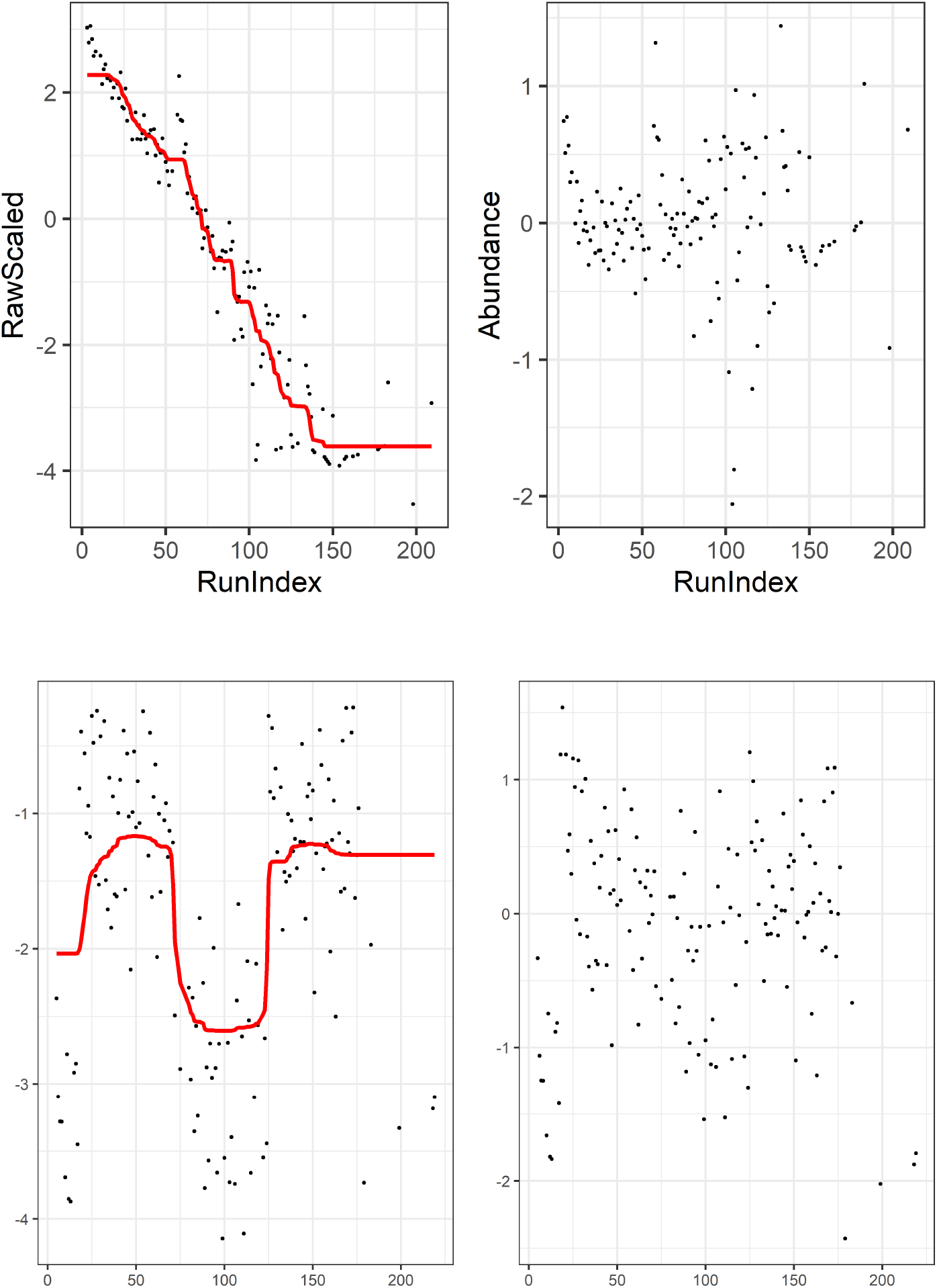
Drift Correction. Two examples of an untargeted metabolite’s drift with a trendline (left), and the resulting abundance after performing drift correction (right)

**eFigure 3:**
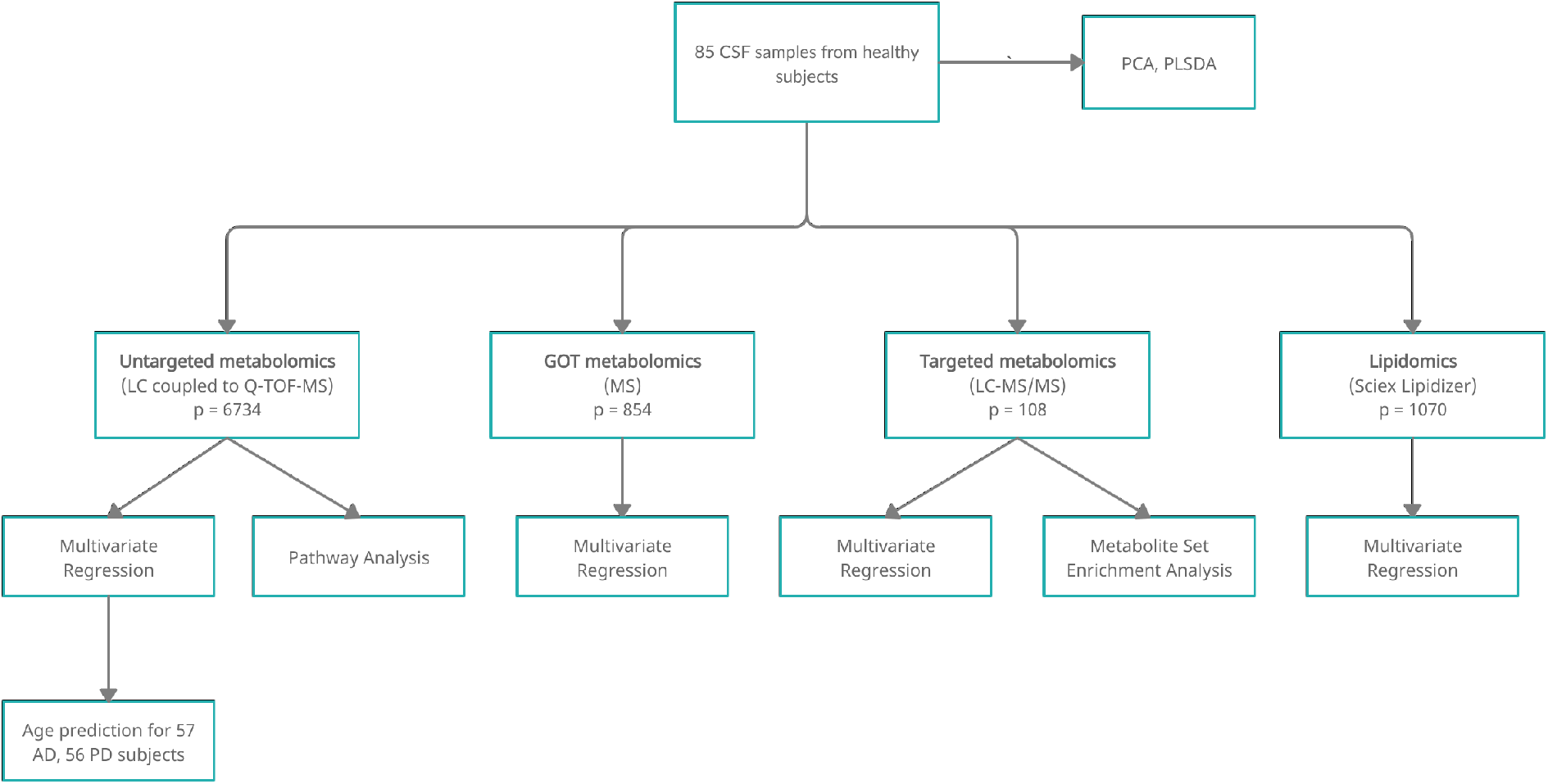
Flowchart outline of analysis. Flowchart outlining the analysis performed on each of the four profiles.

**eFigure 4:**
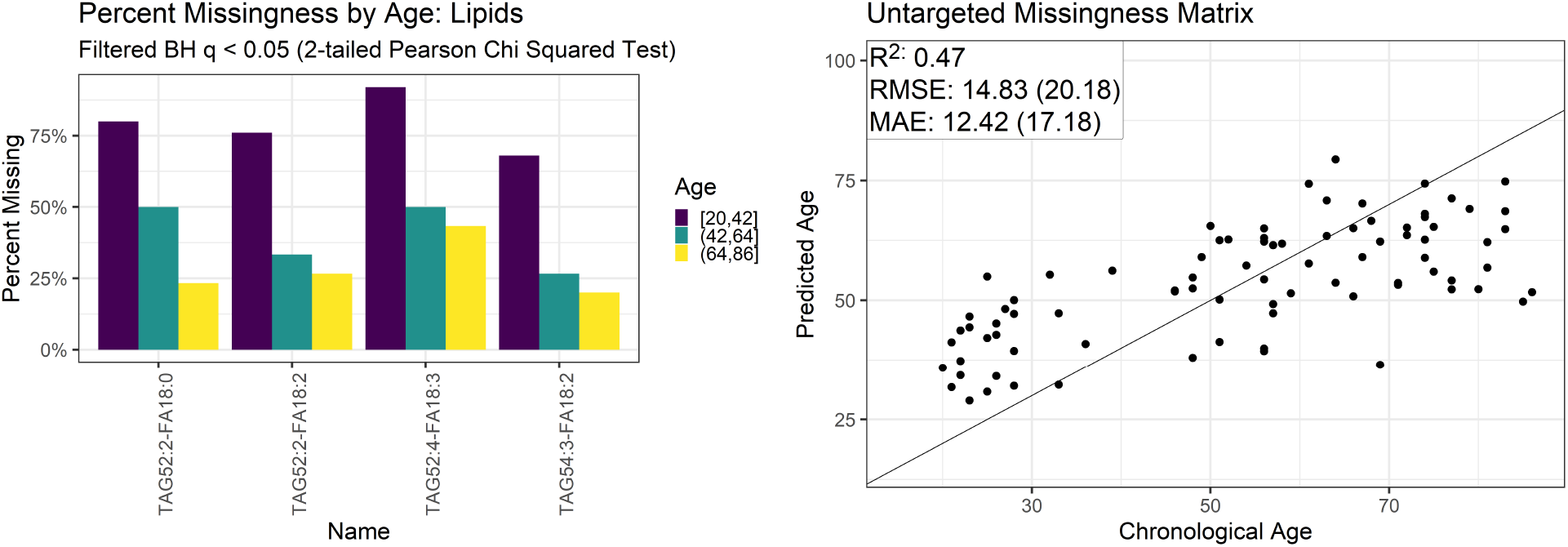
Informative missingness in the lipidome and untargeted metabolome. Informative missingness: Left: Degrees of missingness for lipids that had unequal proportions of missingness at a False Discovery Rate threshold of 0.05. Right: For the untargeted data, we fit elastic net models on age using a matrix of missingness indicators as features, reporting R^2^, Root Mean Squared Error, and Mean Absolute Error. Numbers in brackets represent the performance of the mean model for comparison. A key for the lipid names is provided in eTable 1.

**eFigure 5:**
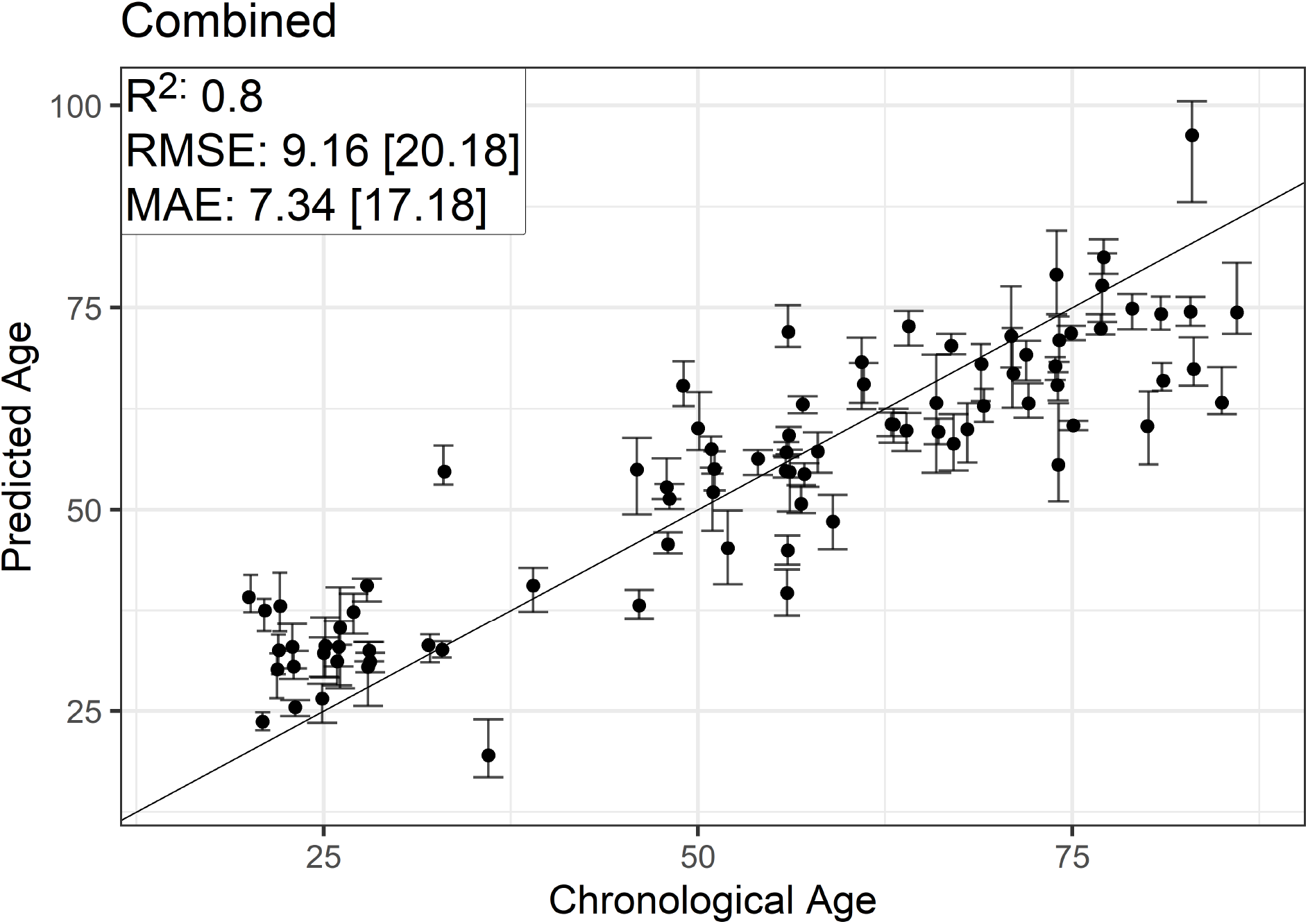
Predictive model for age using combined profiles. Chronological vs Predicted Age for the model combining all four profiles, with R^2^, Root Mean Squared Error (RMSE), and Mean Absolute Error (MAE) reported. Numbers in brackets represent the performance of the mean model for comparison. Points are the average of the predictions for the five imputations to estimate missing values, with the error bars representing the most extreme predicted values from the imputations. We also include the *y* = *x* line. Points above the line correspond to overestimates of a subject’s chronological age, while points below the line correspond to underestimates.

**eTable 2:** Spearman correlation between model predictions. Spearman correlation matrix between the residuals of the models fit using each profile. The residuals are de-trended with respect to age to remove correlation induced by regularization bias. All correlations not involving lipids are significantly non-zero at false discovery rate of 0.05

|  | Targeted | GOT | Untargeted | Lipids |
| --- | --- | --- | --- | --- |
| Targeted | 1 | 0.35 | 0.46 | 0.15 |
| GOT | 0.35 | 1 | 0.54 | 0.1 |
| Untargeted | 0.46 | 0.54 | 1 | 0.23 |
| Lipids | 0.15 | 0.1 | 0.23 | 1 |

**eFigure 6:**
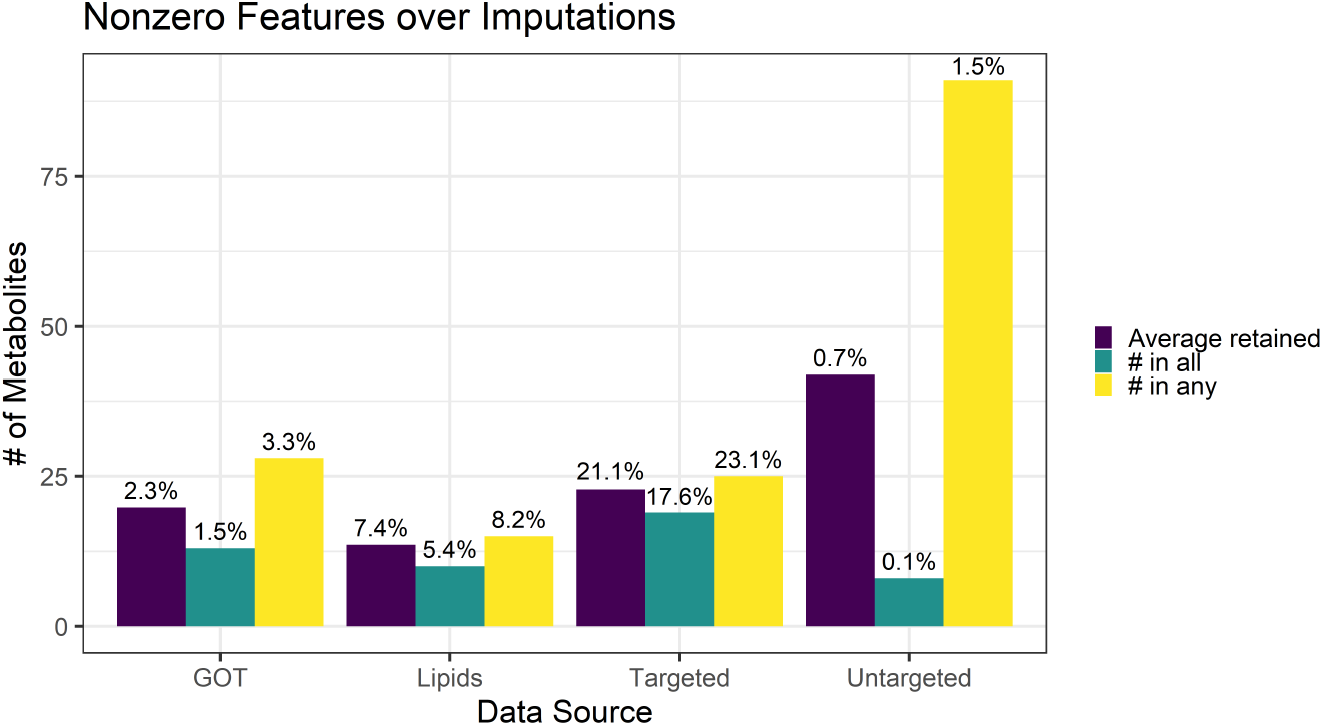
Summary of model size after elastic net regularization. Figure 1: Summary of average model size among the five imputations for each of the four profiles. Percentages above the bars represent the number of metabolites with respect to each of the data sources’ total feature size.

**eFigure 7:**
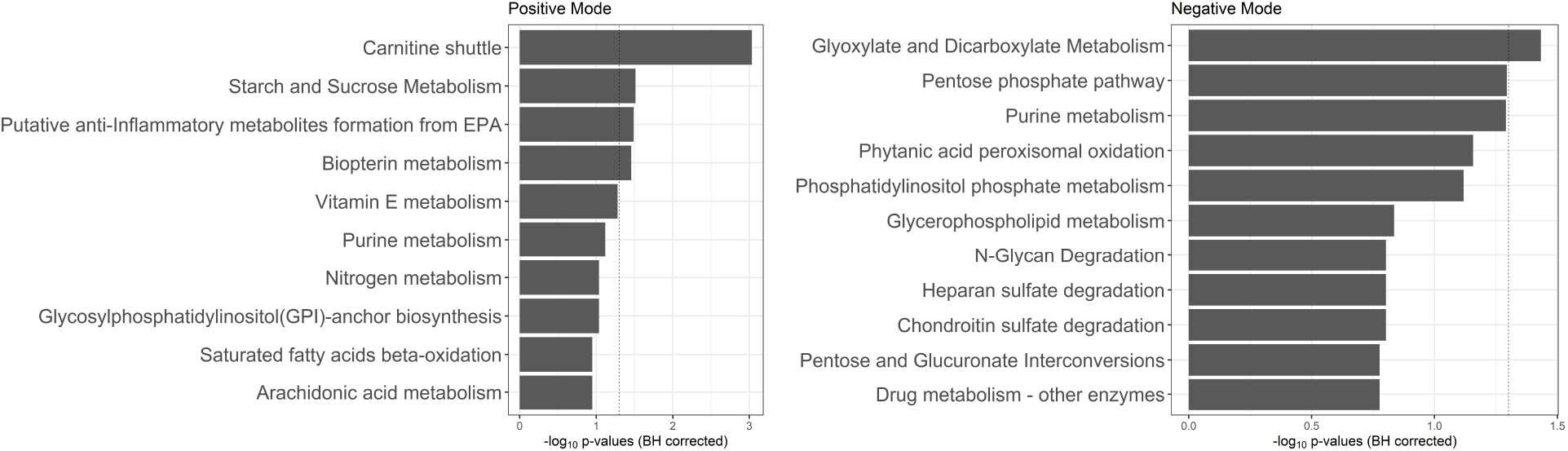
Age-associated pathways identified by Mummichog on the untargeted profile. Pathway Analysis of Mummichog on the untargeted data, sorted by −log_10_ Benjamini Hochberg corrected *p*-values, with the vertical dashed line marking 0.05. Output for the Positive mode is on the left, and Negative mode is on the right

**eTable 3:** MSEA output. Metabolite Set Enrichment Analysis: Output of MSEA on the targeted profile. The input was a list of age-related metabolites from univariate linear analysis. Set are the pre-defined metabolite sets in the SMPDB (left) and CSF library (right). Total is the number of metabolites in these sets, hits is the number of metabolites shared between the SMPDB set and our input list, and expected is the expected number of metabolites shared by random chance, using the cumulative hypergeometric distribution.

| **SMPDB Set** | Total | Expected | Hits |
| --- | --- | --- | --- |
| Tryptophan Metabolism | 8 | 3.86 | 6 |
| Tyrosine Metabolism | 3 | 1.45 | 3 |
| Glutamate Metabolism | 3 | 1.45 | 3 |
| Beta-Alanine Metabolism | 3 | 1.45 | 2 |
| Pyrimidine Metabolism | 3 | 1.45 | 2 |
| Carnitine Synthesis | 3 | 1.45 | 2 |
| Purine Metabolism | 10 | 4.83 | 5 |
| Methionine Metabolism | 8 | 3.86 | 4 |
| Histidine Metabolism | 4 | 1.93 | 2 |
| Betaine Metabolism | 4 | 1.93 | 2 |

| **CSF Library Set** | Total | Expected | Hits |
| --- | --- | --- | --- |
| Parkinson's Disease | 2 | 1.6 | 2 |
| Juvenile Myoclonic Epilepsy | 2 | 1.6 | 2 |
| Post Transurethral Prostatic Resection | 2 | 1.6 | 2 |
| Hydrocephalus | 3 | 2.4 | 2 |
| Aseptic Meningitis | 3 | 2.4 | 2 |
| Lesch-Nyhan Syndrome | 3 | 2.4 | 2 |
| Epilepsy | 2 | 1.6 | 1 |
| Friedreich's Ataxia | 2 | 1.6 | 1 |
| Hereditary Spastic Paraplegia | 2 | 1.6 | 1 |
| Olivopontocerebral Atrophy | 2 | 1.6 | 1 |
| Ataxia (Assorted) | 2 | 1.6 | 1 |
| Friedreich Ataxia | 2 | 1.6 | 1 |
| Heredo-Degenerative Ataxias — Spastic Ataxia | 2 | 1.6 | 1 |
| Hyperphenylalaniemia Due To Guanosine Triphosphate Cyclohydrolase Deficiency | 2 | 1.6 | 1 |
| Meningomyelocoele (Spina Bifida) | 2 | 1.6 | 1 |
| Pyridoxamine 5-Prime-Phosphate Oxidase Deficiency | 2 | 1.6 | 1 |
| Degenerative Disc Disease | 4 | 3.2 | 2 |
| Bacterial Meningitis | 4 | 3.2 | 2 |

### eDiscussion 1: Regularization bias in omic clock models

Figure 3 shows that our models tend to over-predict young subjects and underpredict old subjects, pushing predictions closer to the mean. This behavior could be a result of bias introduced into our model from regularization. Our modeling method of choice, elastic net regression, works by shrinking the size of coefficients down to zero. If enough shrinkage is applied, our model would transform into an intercept-only model, which would place our predictions along a horizontal line at the mean. Thus, when the number of features far exceeds the sample size, the high degree of required regularization can cause the observed prediction bias. To counteract this bias, one can reduce the number of features used in the elastic net model by performing preliminary feature selection. One method, used by Thompson et. al (1), removes covariates with absolute correlation with age < 0.3 prior to each model fit. eFigure 7 shows that this method leads to mixed results on our data, depending on the metabolomic profile. Another method used to mitigate this regularization bias is to perform a variant of what is known as post-selection inference, which involves the following two-stage modeling procedure: the first model is fit on the full set of metabolites and is used as a means of variable selection. The second model is trained only on the metabolites highlighted in the first model, and is used for prediction. The second model is fit using significantly less features, reducing the need for regularization and therefore decreasing the amount of regularization bias. This technique has been used with LASSO regression (a special case of the elastic net), justified under the assumption that the covariates selected by the model are fixed and the signal strength of these covariates is relatively strong (2). These assumptions are likely violated with our data as a result of the small sample size, and the expectation that that our response variable, age, is impacted by many metabolites. eFigure 9 contains two simulations that showcase these principles: one where signal is distributed across a large number of metabolites, and another where signal is concentrated in a small subset of metabolites. Post-selection inference decreases model performance in the first simulation, and slightly improves performance in the second. The epigenetic clock literature suggests that small subsets of predictors are sufficient to achieve a high degree of accuracy, and that there are multiple, almost disjoint subsets that perform comparably. For instance, the 353 feature Horvath model and the 71 feature Hannum model achieve similar predictive accuracy while sharing only 6 CpG sites (3, 4), which suggests the existence of many equally predictive models. The stronger the signal in sparse subsets of predictors, the less likely regularization bias will occur. eFigure 8 shows that the post-selection procedure does not have a large impact on the predictive performance of our models.

**eFigure 8:**
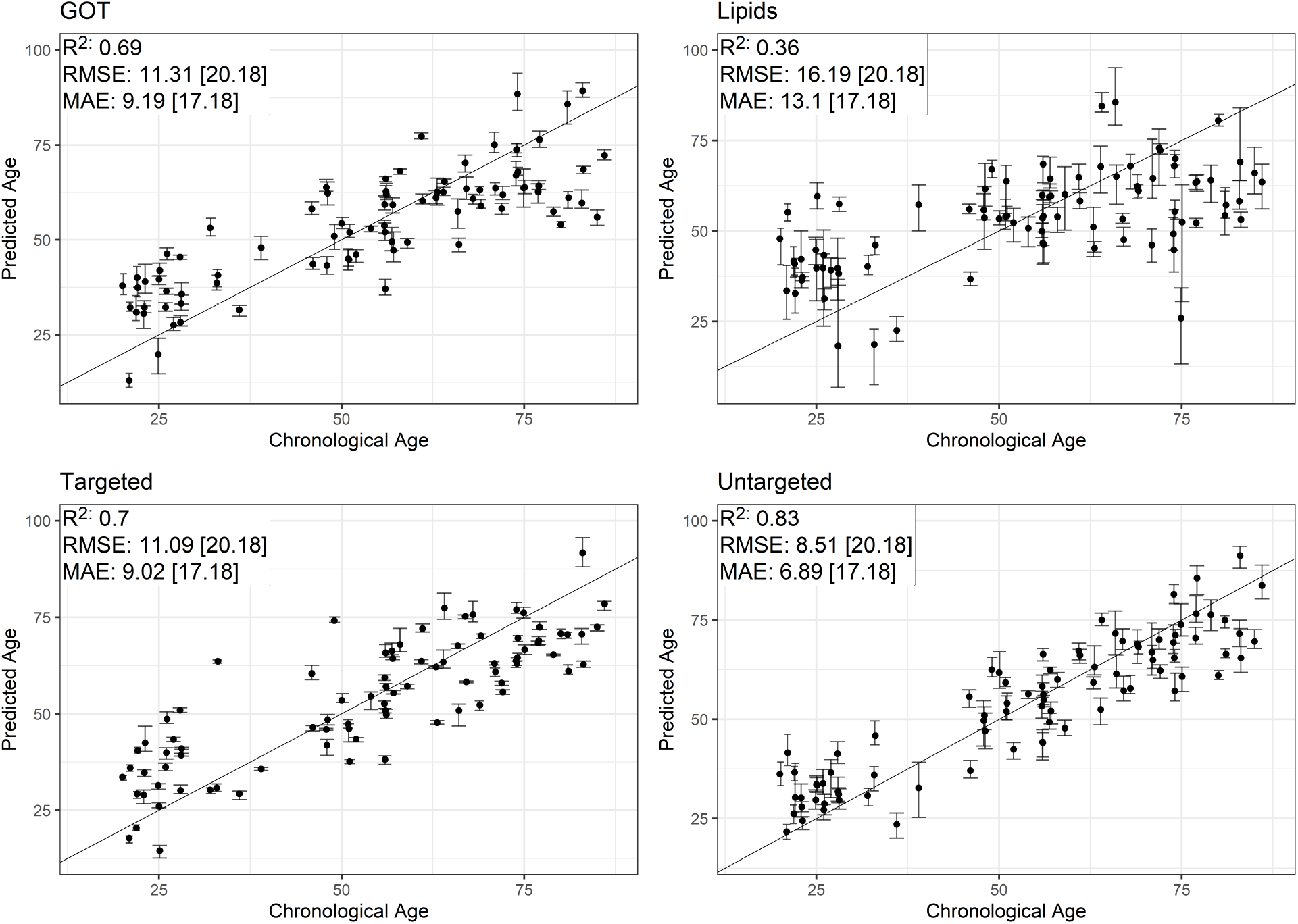
Pre-selection by univariate correlation. Result of pre-selection by absolute correlation > 0.3. Chronological vs predicted age for each dataset, with R^2^, Root Mean Squared Error (RMSE), and Mean Absolute Error (MAE) reported. Numbers in brackets represent the performance of the mean model for comparison. Points are the average of the predictions for the five imputations to estimate missing values, with the error bars representing the most extreme predicted values from the imputations. We also include the *y* = *x* line. Points above the line correspond to overestimates of a subject’s chronological age, while points below the line correspond to underestimates.

**eFigure 9:**
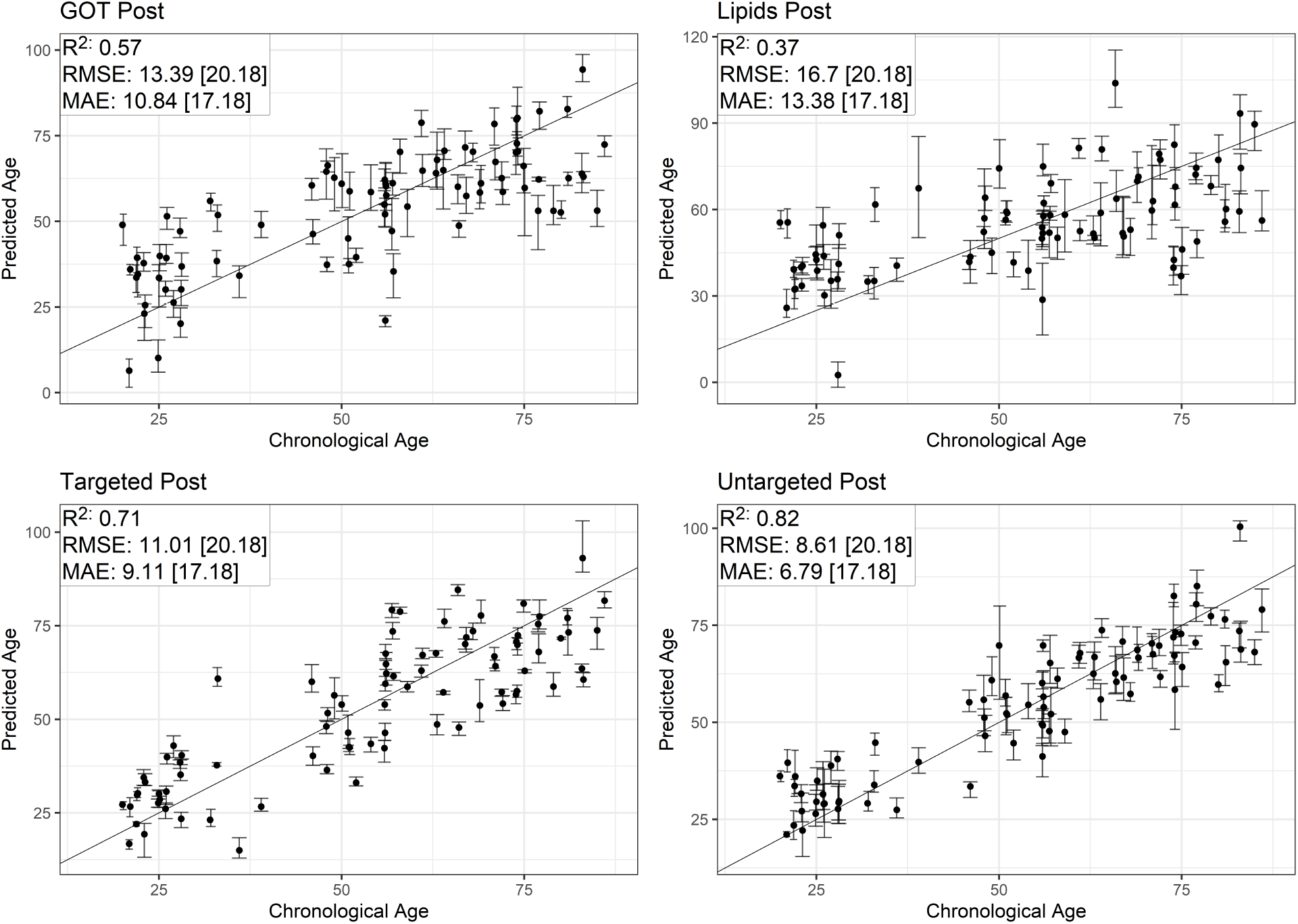
Post-Selection by elastic net. Result of post selection on our profiles. Chronological vs predicted age for each dataset, with R^2^, Root Mean Squared Error (RMSE), and Mean Absolute Error (MAE) reported. Numbers in brackets represent the performance of the mean model for comparison. Points are the average of the predictions for the five imputations to estimate missing values, with the error bars representing the most extreme predicted values from the imputations. We also include the *y* = *x* line. Points above the line correspond to overestimates of a subject’s chronological age, while points below the line correspond to underestimates.

**eFigure 10:**
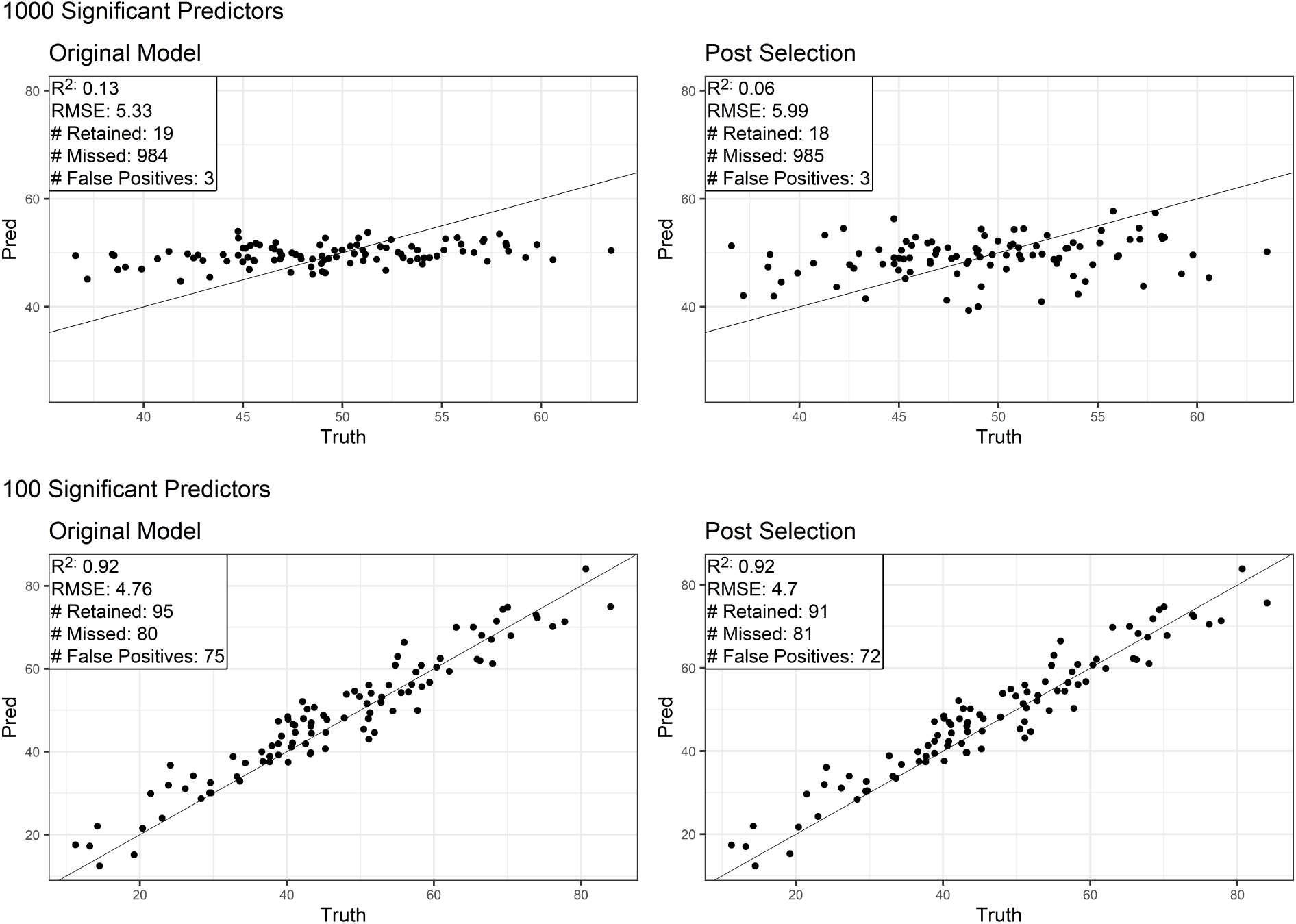
Simulation study highlighting regularization bias. To better understand post selection and its impact on regularization bias, we created a simulated response variable by taking a linear combination of 1000 features in the untargeted dataset and adding a normally distributed error term. Ideally, our regression model would recover the linear combination used to create this variable. However, fitting an elastic net model on half of the subjects and forming a prediction on the second half, we found that the model selected only 16 of the 1000 truly significant features, likely contributing to the pattern of over predicting early and under predicting late. In this case, we see that fitting a post-selection model on the 19 nonzero features is unhelpful, as it is fit on very few of the significant metabolites, and even increases error. On the other hand, consider the setup above, but with the modification that the simulated response variable is a linear combination of only 100 metabolites (with similar amounts of total signal, so that the signal to noise ratio in the significant metabolites is comparatively larger). In this instance, the elastic net model has low error, but selects only 20 of the 100 significant metabolites. In this instance, the signal from these metabolites is strong enough to make accurate predictions, so post-selection modeling exhibits little effect on the prediction. Models fit using the untargeted dataset and simulated response variables to show the impact of regularization bias and post selection inference. The first row shows that regularization bias can occur when the signal to noise ratio for each metabolite is small, and that post selection procedures can reduce model performance by increasing variance. The second row shows that regularization bias is not likely to occur when the true model is sparse, and that post selection procedures are not likely to have significant impact on the model.

**eFigure 11:**
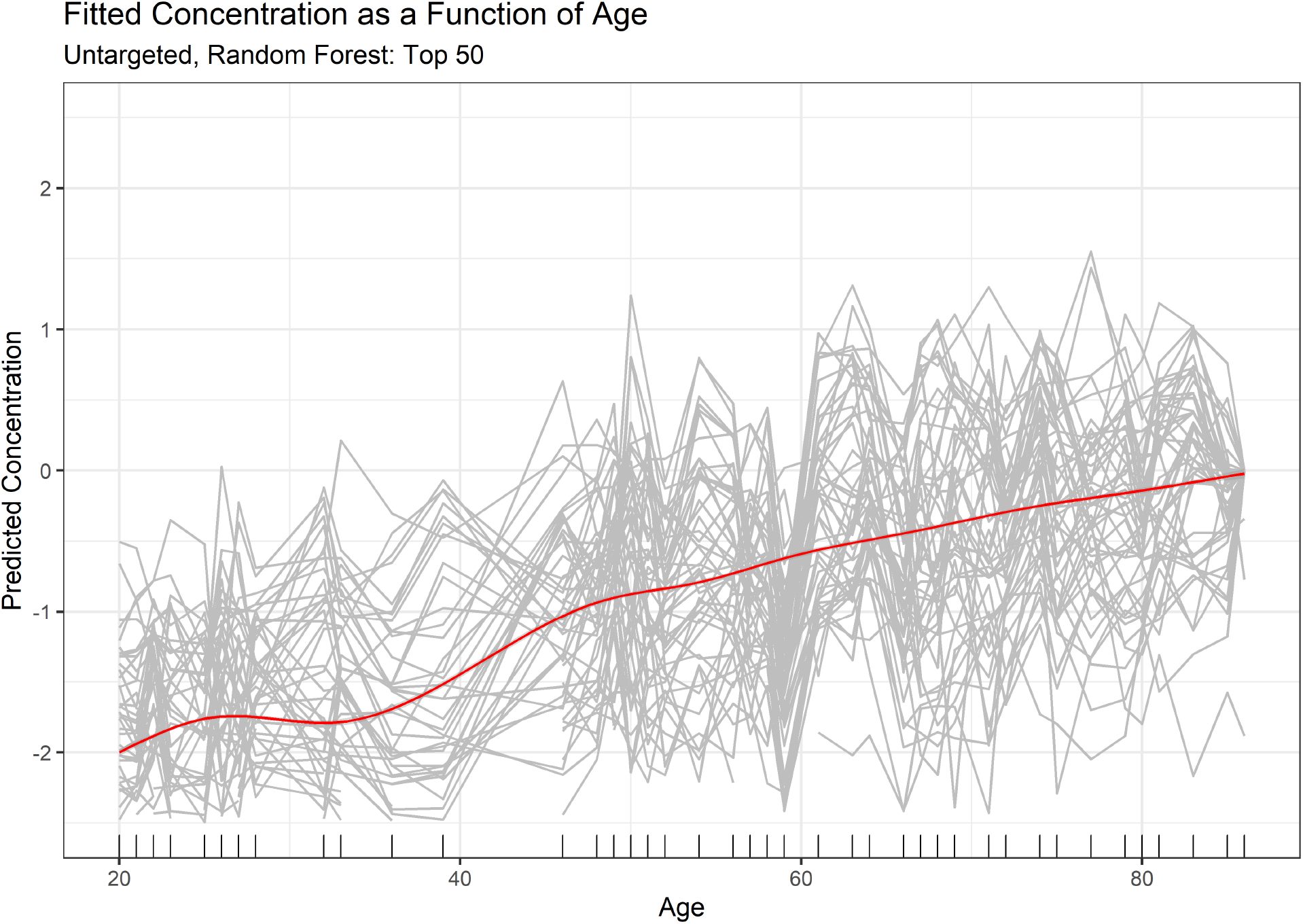
Predicting metabolite concentration using age. Fitting random forests on each of the metabolite concentrations as a function of age, and displaying the top fifty metabolites in terms of variance explained. The gray lines are fitted concentrations for individual metabolites, while the red line is an average.

## Notes

### Competing Interest Statement

The authors have declared no competing interest.

